# IRE1β negatively regulates IRE1α signaling in response to endoplasmic reticulum stress

**DOI:** 10.1101/586305

**Authors:** Michael J. Grey, Eva Cloots, Mariska S. Simpson, Nicole LeDuc, Yevgeniy V. Serebrenik, Heidi De Luca, Delphine De Sutter, Phi Luong, Jay R. Thiagarajah, Adrienne W. Paton, James C. Paton, Markus A. Seeliger, Sven Eyckerman, Sophie Janssens, Wayne I. Lencer

**Author notes:** These authors contributed equally to this work (Michael J. Grey, Eva Cloots, Mariska S. Simpson). Corresponding author: Wayne I. Lencer, Division of Gastroenterology, Hepatology, and Nutrition, Boston Children’s Hospital, 300 Longwood Avenue, Enders 609, Boston MA 02115.

## Abstract

IRE1β is an ER stress sensor uniquely expressed in epithelial cells lining mucosal surfaces. Here, we show that intestinal epithelial cells expressing IRE1β have an attenuated response to ER stress. IRE1β assembles with and blocks activation of the closely related and most evolutionarily ancient stress-sensor IRE1α to suppress stress-induced xbp1 splicing, a key mediator of the unfolded protein response. In comparison, IRE1β has weak xbp1 splicing activity, largely explained by a non-conserved amino acid in the kinase domain that impairs its phosphorylation and restricts oligomerization. This enables IRE1β to act as a dominant negative suppressor of IRE1α. The inhibitory effect is amplified in cells by disrupting an XBP1-dependent feedback loop regulating stress-induced expression of IRE1α. Thus IRE1β functions to negatively regulate IRE1α signaling, perhaps enabling intestinal epithelial cells to manage the response to chronic stress stimuli at the host-environment interface.

## INTRODUCTION

All mammalian cell types have three sensors in the endoplasmic reticulum - IRE1α, ATF6, and PERK - which detect imbalances in protein folding and trigger an integrated set of signaling pathways to restore normal proteostasis. This is called the unfolded protein response (UPR). If protein folding in the ER remains unresolved, prolonged UPR signaling induces cell death (Hetz and Papa, 2018; Walter and Ron, 2011). Epithelial cells lining the intestine and other mucosal surfaces that interface with the environment are unique in that they express an additional ER stress sensor called IRE1β (*ern2* gene) (Bertolotti et al., 2001; Iwawaki et al., 2001; Martino et al., 2013; Tsuru et al., 2013; Wang et al., 1998). IRE1β is a close paralogue of the ubiquitously expressed IRE1α (Tirasophon et al., 1998). Both are dual kinase/endonucleases that splice xbp1 mRNA to produce the transcription factor XBP1s, which functions to induce the UPR (Calfon et al., 2002; Lee et al., 2002; Yoshida et al., 2001). Both IRE1α and IRE1β can also degrade other mRNA sequences targeted to the ER for translation - termed Regulated IRE1-Dependent Decay of mRNA, or RIDD (Hollien et al., 2009; Hollien and Weissman, 2006; Imagawa et al., 2008; Iwawaki et al., 2001; Tsuru et al., 2013) - including for IRE1α the ability to autoregulate its own expression by degrading its own mRNA (Tirasophon et al., 2000). Yet, despite the high degree of sequence homology between the two molecules, IRE1β and IRE1α appear to have distinct enzymatic activities, and how IRE1β functions in the ER stress response remains inconclusively defined. In cell culture, some studies show that IRE1β can sense ER stress and activate the UPR by splicing xbp1 transcripts (Tirasophon et al., 2000; Wang et al., 1998). But other reports suggest it is less effective than IRE1α at splicing xbp1 and signals through other mechanisms to mitigate ER stress (Imagawa et al., 2008; Iwawaki et al., 2001).

In vivo, under normal physiologic conditions, the intestine and colon of mice lacking IRE1β (IRE1β^-/-^) show evidence for an elevated UPR compared to wild type controls, including increased levels of spliced xbp1 transcript indicative of IRE1α activation (Bertolotti et al., 2001; Tschurtschenthaler et al., 2017; Tsuru et al., 2013). The phenotype suggests that IRE1β may function to suppress IRE1α activity, and perhaps other elements of the UPR. Such a role for IRE1β in diminishing ER stress in the intestine was most recently implicated in mice conditionally lacking both the IRE1α substrate XBP1 and the autophagy factor ATG16L1 (Tschurtschenthaler et al., 2017). At the molecular level, activation of IRE1α by ER stress appears to require homo-oligomerization and autophosphorylation (Bertolotti et al., 2000; Li et al., 2010). Given the close homology between the two proteins, we became interested in testing the hypothesis that IRE1β may modulate the UPR by interacting and assembling directly with IRE1α. We examined IRE1β function in HEK293 cells and in vitro using purified proteins. Our cell and biochemical data show that IRE1β restricts ER stress-induced IRE1α endonuclease activity, as assessed by xbp1 splicing, and it reverses the increases in IRE1α and XBP1 expression expected for the canonical UPR. We define structural features of the IRE1β kinase domain that explain these effects and enable IRE1β to act as a direct and dominant-negative suppressor of IRE1α oligomerization and signaling. This activity appears to have been evolutionarily conserved in epithelial cells lining the mucosa of vertebrates, perhaps, as proposed before (Bertolotti et al., 2001), to dampen amplified ER stress responses inherent to the host-environment interface.

## RESULTS

### IRE1β suppresses IRE1α signaling in response to ER stress

Previous studies suggest that IRE1β restricts IRE1α signaling in vivo under normal homeostatic conditions (Bertolotti et al., 2001; Tschurtschenthaler et al., 2017; Tsuru et al., 2013). To test this idea, we compared IRE1α signaling in polarized human intestinal epithelial cell lines expressing different levels of IRE1β. The human intestinal T84 cell line expresses IRE1β but the human intestinal Caco2 cell line does not (Figure 1A, left panel). Both cell lines express IRE1α at steady state, though Caco2 cells express more (Figure 1A, middle panel, untreated). When treated with thapsigargin to induce ER stress, Caco2 cells responded with a robust increase in spliced xbp1 transcripts indicative of IRE1α activation (Figure 1A, right panel). They also responded with a 4-fold increase in IRE1α mRNA expression (Figure 1A, middle panel), an expected outcome for induction of the UPR. In contrast, T84 cells, which expresses IRE1β, responded to ER stress with significantly lower levels of spliced xbp1 and no detectable increase in *ern1* mRNA expression (Figure 1A, middle and right panels). The same results were obtained using primary intestinal colonoids prepared from wild type (IRE1β^+/+^) or IRE1β^-/-^ mice (Figure 1B). In this case, ER stress was induced using subtilase cytotoxin (SubAB) (Paton et al., 2006; Paton et al., 2004) as validated in this model before (Heijmans et al., 2013; Wielenga et al., 2015). Similar to the Caco2 and T84 cell lines treated with thapsigargin, lower levels of xbp1 splicing was found after SubAB treatment of colonoids prepared from IRE1β^+/+^ mice as compared to those prepared from IRE1β^-/-^ mice. Thus, IRE1β expression correlates with reduced IRE1α activity (as measured by xbp1 splicing) and IRE1α gene expression, both in response to the induction of ER stress.

**Figure 1.**
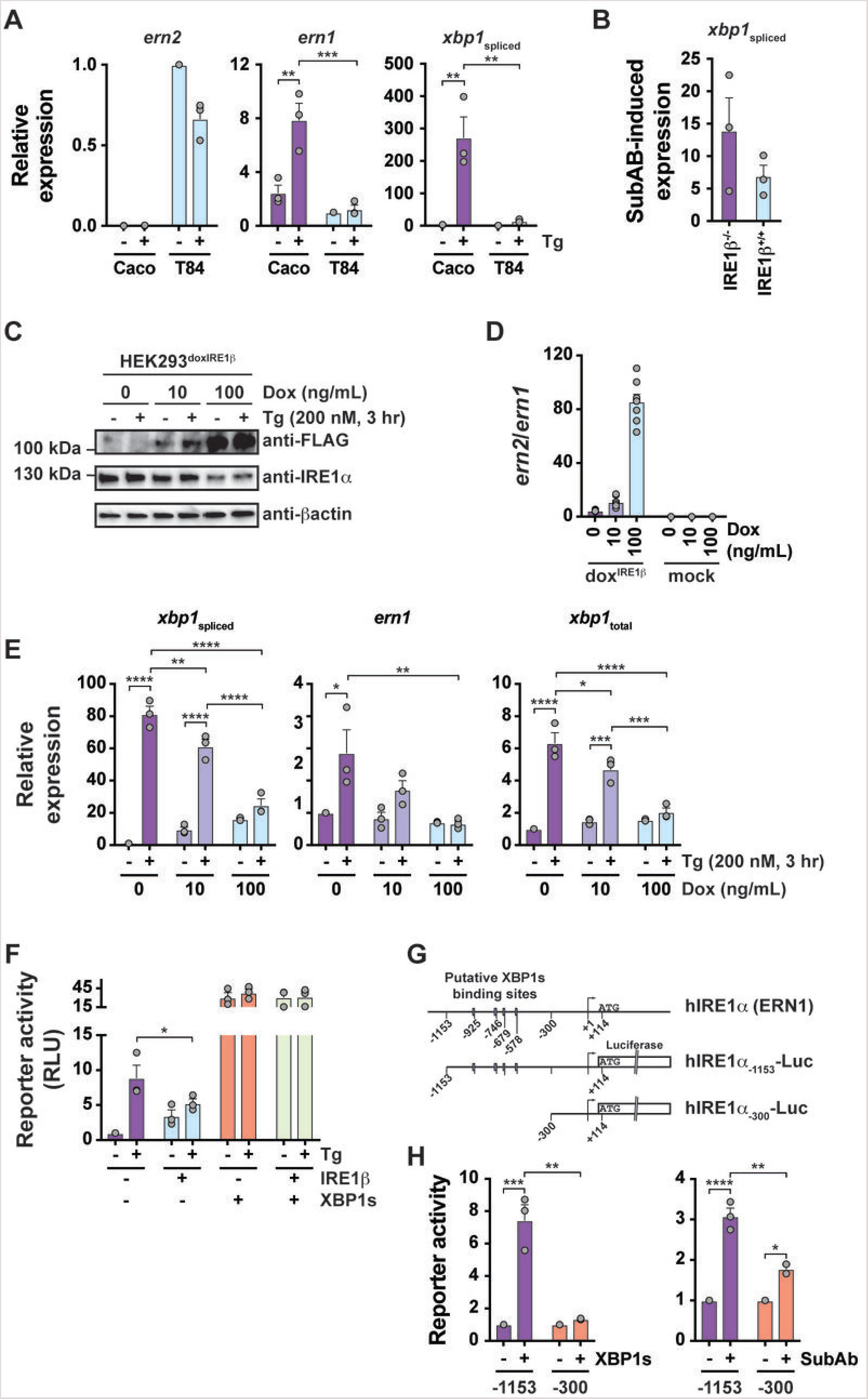
IRE1β suppresses IRE1α signaling in response to ER stress. (A) Expression of *ern2, ern1*, and spliced xbp1 transcripts were assayed by qPCR for polarized Caco2 and T84 monolayers treated without or with 3 μM thapsigargin (Tg) for 2 hr. Bars represent mean ± SEM for 3 independent experiments. (B) Spliced xbp1 transcript was assayed by qPCR for IRE1β^+/+^ and IRE1β^-/-^ mouse primary colonoids treated with 100 ng/mL subtilase cytotoxin (SubAB) for 24 hrs. Expression is plotted relative to colonoids treated with an equivalent dose of an inactive mutant toxin SubA_A272_B. Bars represent mean ± SEM for 3 independent experiments. (C-E) IRE1β expression was induced in HEK293^doxIRE1β^ cell line with indicated concentration of doxycycline (Dox) for 24 hr. Cells were stimulated with 200 nM thapsigargin for 3 hr and assayed for (A) IRE1β-FLAG and IRE1α protein expression by western blot, (D) relative *ern2*/*ern1* transcript levels by qPCR, and (E) spliced *xbp1, ern1*, and total *xbp1* transcripts by qPCR. The blot in (C) is representative of 3 independent experiments. The bars in (D) and (E) represent mean ± SEM for 8 and 3 independent experiments, respectively. (F) HEK293 cells were co-transfected with UPRE-luciferase reporter and indicated plasmids, treated with thapsigargin for 8 hr, and assayed for luciferase activity. Bars represent mean ± SEM for 3 independent experiments. (G) Schematic of human IRE1α gene (*ern1*) promoter region and luciferase reporter constructs. Putative XBP1s binding sites are indicated as boxes. (H) IRE1α-Luc reporter activity in HEK293 cells (left) co-transfected with either control vector or XBP1s expression vector or (right) treated with SubA_A272_B or SubAB (100 ng/mL) for 8 hr. Bars represent mean ± SEM for 3 independent experiments.

To test if IRE1β was sufficient to suppress IRE1α signaling, we generated a HEK293 cell model with doxycycline (Dox)-inducible expression of FLAG-tagged IRE1β (HEK293^doxIRE1β^). Treatment of these cells with Dox for 24 hr induced IRE1β expression in a dose-dependent manner (Figure 1C, anti-FLAG blot). At the transcript level, this corresponded to approximately 10- and 100-fold higher levels of IRE1β (*ern2*) relative to the endogenous expression of IRE1α (*ern1*), after treatments with 10 or 100 ng/mL Dox, respectively (Figure 1D). This is similar to the expression levels observed for IRE1β and IRE1α in the normal mouse small intestine (Haber et al., 2017). In goblet cells, for example, where IRE1β is most highly expressed, the *ern2*:*ern1* mRNA transcript ratio as measured in single cells ranged from 2:1 to 384:1 with a median value of 22:1 (*ern2:ern1*, Supplemental Figure 1A). Thus, the HEK293^doxIRE1β^ cell model approximates physiologic levels of IRE1β expression compared to IRE1α.

In the absence of IRE1β (no Dox), Tg treatment (3 hr) of HEK293^doxIRE1β^ cells caused increased IRE1α (*ern1*) and *xbp1* expression, and IRE1α activation as measured by spliced *xbp1* transcript (*xbp1*_spliced_) (Figures 1E: no dox, compare ± Tg). Similar results were obtained with HEK293 cells lacking the IRE1β lentiviral transgene (HEK293^mock^, Supplemental Figure S1B). Thus, in the absence of IRE1β expression (no dox), the HEK293^doxIRE1β^ model exhibits key features of IRE1α activation associated with the conventional UPR. When HEK293^doxIRE1β^ cells were treated with 100 ng/mL Dox for 24 hr to induce IRE1β expression, *xbp1* splicing in response to Tg was significantly reduced, consistent with inhibition of IRE1α activation by IRE1β expression (Figure 1E, left panel). There was also significant reduction in xbp1 splicing at the lower level of Dox treatment compared to no Dox, suggesting a causal effect of IRE1β expression. Similar results were obtained in cells treated with Tg for shorter (1.5 hr) or longer time points (6 hr, Supplemental Figure S1C), implicating a rapid or pre-existing effect of IRE1β on IRE1α that is sustained in the presence of ER stress. There was no effect of Dox treatment on baseline or stress-induced levels of spliced xbp1 transcripts in HEK293^mock^ cells (Supplemental Figure S1B). In the HEK293^doxIRE1β^ model, there was also significant reduction in Tg-induced *ern1* and *xbp1* transcript levels (Figure 1E, middle and right panels), consistent with broader suppression of the UPR to ER stress. Overexpression of IRE1β by transient transfection yielded a similar phenotype, with significantly reduced stress-induced spliced *xbp1* and *ern1* transcript levels (Supplemental Figure S1D).

The apparent inhibitory effect of IRE1β also extended downstream of IRE1α to expression from XBP1 dependent promoters. In HEK293 cells containing an unfolded protein response element (UPRE) luciferase reporter, XBP1s protein (a transcription factor) drives luciferase expression from five UPRE sites (Wang et al., 2000). In control cells lacking IRE1β, Tg treatment induced robust expression of the luciferase reporter as expected for the UPR (Figure 1F, purple bars). In cells expressing IRE1β, reduced levels of XBP1s protein (Supplemental Figure S1E) correlated with significantly attenuated reporter activity (Figure 1F, blue bars). Overexpression of XBP1s protein rescued UPRE reporter activity (Figure 1F, orange and green bars) indicating that IRE1β acts upstream of XBP1s. These results suggested an explanation for how IRE1β may suppress IRE1α expression by inhibiting XBP1 transcriptional activity (Figures 1C and 1E). To test this, we analyzed the human *ern1* gene sequence and found four XBP1s binding sites in the distal promoter region 300-1200 bp upstream of the transcription start site (Figure 1G). Luciferase reporter constructs containing these sites were activated both by overexpression of XBP1s and ER stress stimulation (Figure 1H). Thus IRE1α can regulate its own expression, at least in part, by producing XBP1s in response to ER stress. Altogether, these studies show the HEK293^doxIRE1β^ model recapitulates the in vivo phenotypes observed in IRE1β^-/-^ mice and implicate IRE1β in the suppression of IRE1α expression and signaling in the UPR.

### IRE1β blocks IRE1α activation and oligomer assembly

As the UPR is associated with IRE1α oligomer formation (Bertolotti et al., 2000; Li et al., 2010), we tested if IRE1β affects IRE1α oligomerization. These studies were done using HEK293 cells overexpressing IRE1β. ER stress was induced using the N-glycosylation inhibitor tunicamycin. The anticipated oligomerization and shift of endogenous IRE1α to higher molecular weight species in tunicamycin-treated cells was assessed by gel filtration chromatography of detergent cell lysates and immunoblot for endogenous IRE1α. In mock-transfected cells lacking IRE1β, tunicamycin caused IRE1α to elute in earlier fractions compared to tunicamycin untreated cells (Figure 2A, upper two gels). This shift to higher apparent molecular weight is presumed to reflect IRE1α oligomer formation. Transfection with IRE1β, on its own, also caused IRE1α to elute in higher molecular weight fractions, though to a far lesser extent than that caused by ER stress in cells lacking IRE1β. The shift to higher molecular weight presumably represents a degree of activation of IRE1α, but this was not further enhanced by tunicamycin treatment (Figure 2A, bottom two gels). Similar results were obtained using the IRE1β-inducible HEK293^doxIRE1β^ model treated with thapsigargin to induce ER stress (Figure 2B). These results suggest that expression of IRE1β impairs IRE1α oligomerization (and thus activation) following ER stress, although an alternative interpretation could be that IRE1β over-expression on its own induced a state of ER stress obfuscating the effect of tunicamycin or thapsigargin treatment.

**Figure 2.**
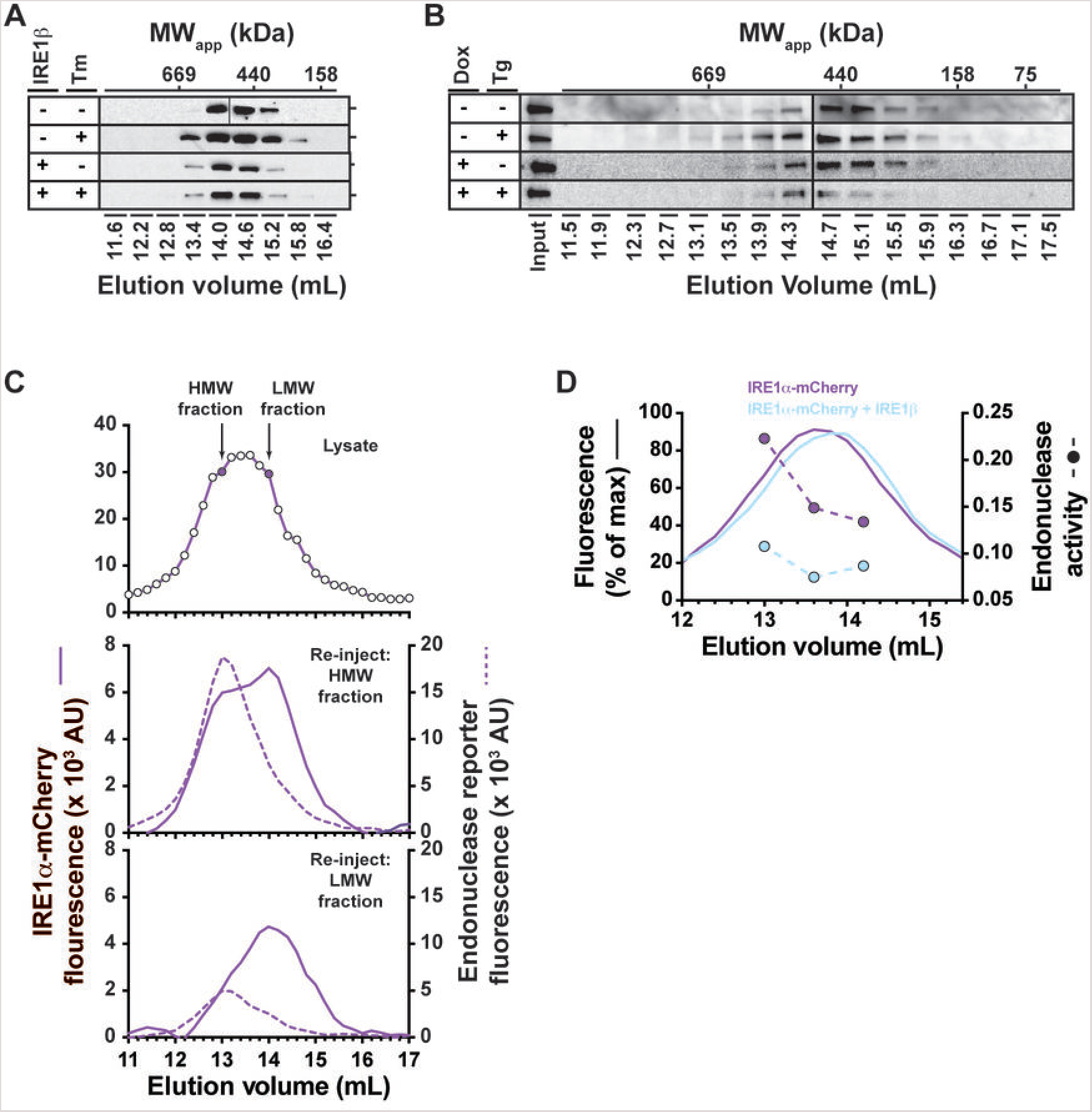
IRE1β blocks IRE1α activation and oligomer assembly. (A) Mock-transfected and IRE1β-transfected HEK293T cells were treated with tunicamycin (500 ng/mL for 4 hr), solubilized with dodecylmaltoside lysis buffer, and fractionated by gel filtration chromatography. IRE1α elution was assayed by western blot with anti-IRE1α antibody. (B) IRE1β expression was induced in HEK293^doxIRE1β^ cell line with 100 ng/mL doxycycline (Dox) for 24 hr. Cells were stimulated with 200 nM thapsigargin for 6 hr and analyzed by gel filtration as in (A). (C, top panel) HEK293 cells expressing IRE1α-mCherry were solubilized with dodecylmaltoside and fractionated by gel filtration chromatography. IRE1α elution was assayed by mCherry fluorescence. The “high molecular weight” (HMW) and “low molecular weight” (LMW) fractions (indicated by filled symbols in top panel) were re-injected on gel filtration. mCherry fluorescence (solid lines) and endonuclease activity (dashed lines) are shown for HMW (middle panel) and LMW (bottom panel) samples. (D) HEK293 cells expressing IRE1α-mCherry alone or IRE1α-mCherry + IRE1β-MycHis were solubilized with dodecylmaltoside and fractionated by gel filtration chromatography. Elution of IRE1α was assayed by mCherry fluorescence (solid lines). *In vitro* endonuclease activity was measured for individual fractions with 1000 nM xbp1 reporter substrate. Endonuclease activity (symbols, dashed lines) is plotted as AU min^-1^ mCherry^-1^. Chromatograms are representative of 2 independent experiments.

Given that uncertainty, we tested the idea another way using HEK293 cells expressing mCherry-tagged IRE1α (IRE1α-mCherry) with and without IRE1β co-expression at steady state. In this case, over-expression of IRE1α-mCherry was presumed to induce a state of ER stress, as shown previously in other systems (Li et al., 2010; Tirasophon et al., 1998). Detergent cell lysates were assayed by gel filtration chromatography for IRE1α-mCherry oligomer formation using direct fluorescence. When expressed on its own, IRE1α-mCherry eluted as a mix of species with elution of a high molecular weight species around 13.0 mL and a lower molecular weight species 14.0 mL (the profiles varied between a broad peak centered at 13.6 mL as shown in Figure 2C (upper panel) or as more prominent high and low molecular weight peaks as shown in Figure 4E). This is similar to the elution profile for endogenous IRE1α in lysates from stress-stimulated cells (Figures 2A and 2B). When the higher (13.0 mL fraction) or lower (14.0 mL fraction) molecular weight species were re-injected on gel filtration, both forms were again obtained (Figure 2C, middle and bottom panels), suggesting that the higher and lower molecular weight species in cell lysates were in thermodynamic equilibrium and represent distinct oligomeric forms (tetramers versus dimers, respectively; discussed further below). Endonuclease activity was also measured on each fraction using an in vitro assay for cleavage of a model xbp1-reporter substrate (Wiseman et al., 2010). Notably, the highest levels of endonuclease activity co-eluted with the higher molecular weight species at 13.0 mL, suggesting that this represents the active, stress-stimulated form. In HEK293 cells co-expressing IRE1β, the elution profile for IRE1α-mCherry was shifted subtly but reproducibly towards lower apparent molecular weight (Figure 2D, light blue trace compared to purple trace). There was also a concomitant reduction in in vitro endonuclease activity for the higher molecular weight species (Figure 2D, light blue symbols). These results are consistent with IRE1β blocking (or altering) IRE1α oligomer formation and endonuclease activity.

### IRE1β interacts directly with IRE1α to inhibit xbp1-splicing

To explain how IRE1β suppresses IRE1α, we first asked if the molecules’ enzymatic activities were involved. We prepared IRE1β expression constructs with inactivating point mutation in its kinase domain (IRE1βK574A), with deletion of the endonuclease domain (IRE1β(Δ783-925)), and with deletion of both endonuclease and kinase domains combined (IRE1β(Δ453-925)) (Figure 3A, schematic). Full length IRE1β or mutant IRE1β constructs were co-expressed along with a luciferase reporter for xbp1-splicing (Iwawaki and Akai, 2006) in HEK293 cells that express endogenous IRE1α. All IRE1β constructs were expressed at similar levels (Figure 3A, immunoblots). ER stress was induced using thapsigargin (Tg) or SubAB, and using the toxin’s enzymatically-inactive mutant (SubA_A272_B) as control. SubA_A272_B enters the ER of host cells like SubAB but cannot induce ER stress (Paton et al., 2006; Paton et al., 2004).

**Figure 3.**
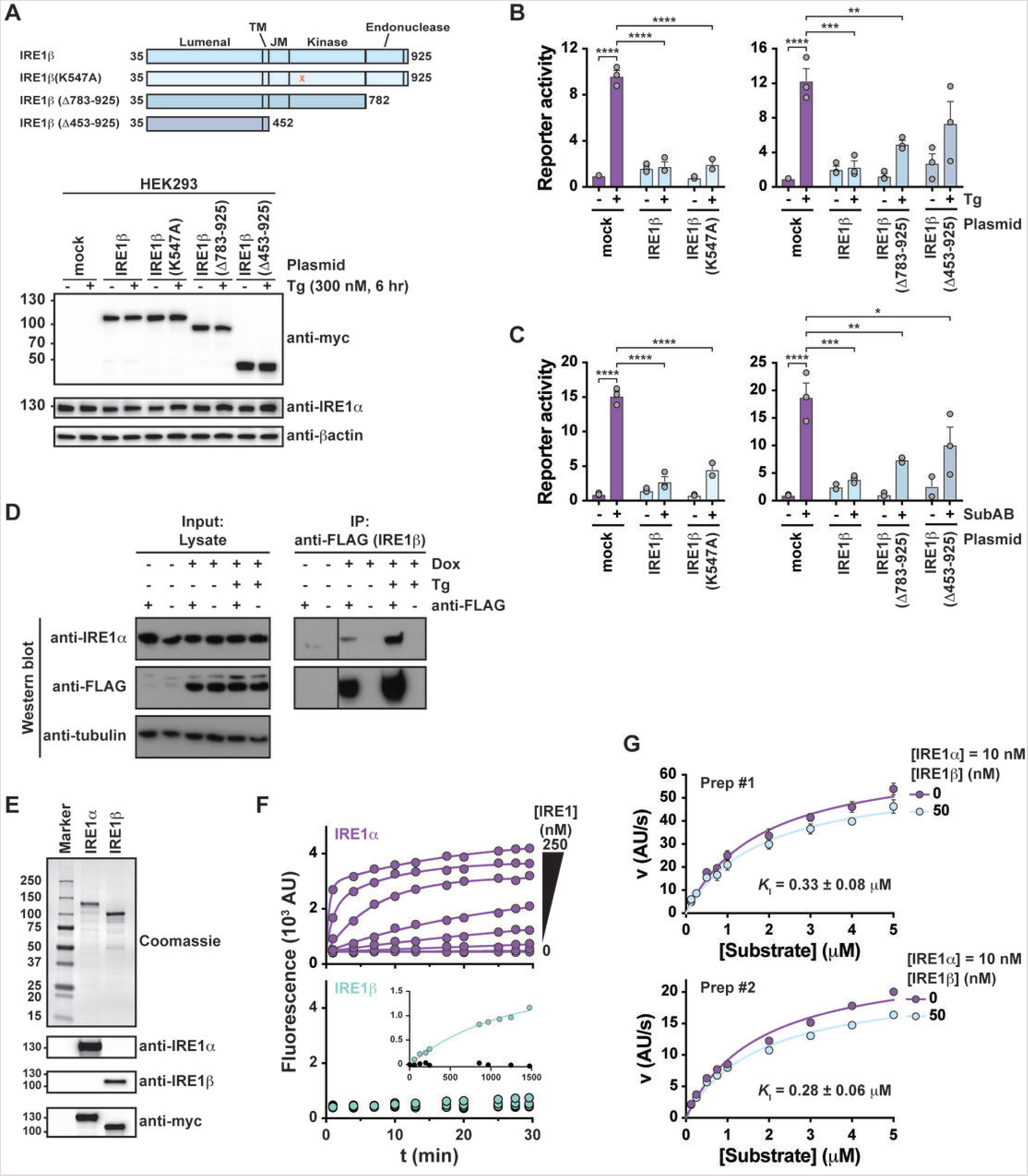
IRE1β interacts directly with IRE1α to inhibit xbp1 splicing. (A) Schematic of IRE1β constructs. All constructs have C-terminal myc tag. Constructs were overexpressed in HEK293 cells and expression assayed by western blot with anti-myc antibody. (B-C) Xbp1 splicing activity was measured using a luciferase reporter in HEK293 cells co-transfected with indicated IRE1β expression plasmids and stimulated with either (B) thapsigargin (Tg, 3 μM) or (C) SubAB (100 ng/mL) for 8 hr at 37 °C. Luciferase activity was measured using Bright Glo luminescence assay and is plotted relative to mock-transfected, control-treated cells. Bars represent mean ± SEM for 3 independent experiments. (D) IRE1β expression was induced in HEK293^doxIRE1β^ cell line with 100 ng/mL doxycycline (Dox) for 24 hr. Cells were treated with thapsigargin, lysed in E1A buffer, incubated with biotinylated anti-FLAG antibody, and immunoprecipitated with streptavidin-coated magnetic beads. Lysate and immunoprecipitated samples were assayed by western blot with anti-IRE1α and anti-FLAG antibodies. Blots are representative of 3 independent experiments. (E) Samples of affinity-purified full-length IRE1α and IRE1β were separated on SDS-PAGE and stained with Coomassie blue or assayed by western blot with anti-IRE1α, anti-IRE1β, or antimyc antibodies. (F) In vitro endonuclease activity was assayed by monitoring cleavage of fluorescent reporter substrate (10 nM) over time for indicated concentrations of (top, purple) IRE1α or (bottom) IRE1β. (Bottom, inset) Endonuclease activity monitored over 24 hr for 250 nM IRE1β (light green) or buffer control (black circles). (G) Steady state kinetics were assayed by measuring progress curves for 10 nM IRE1α or 10 nM IRE1α + 50 nM IRE1β as a function of reporter substrate concentration. Kinetic data are plotted as initial reaction velocity versus substrate concentration. Data were measured for two independent preparations of purified protein, with symbols representing mean ± SEM for 3 independent measures for Prep#1 or values from a single measurement for Prep#2. Solid lines represent best fit of a non-competitive inhibitor model to the kinetic data.

Mock-transfected cells at baseline displayed low levels of xbp1-splicing reporter activity that was significantly increased by treatment with Tg (Figure 3B) or SubAB (Figure 3C). The robust stress-induced xbp1-splicing seen in mock-transfected HEK293 cells was nearly completely reversed in cells overexpressing full length IRE1β. The same result was obtained in cells overexpressing the IRE1β kinase dead mutant IRE1β(K547A), indicating that kinase activity is not required for IRE1β to inhibit IRE1α. Similarly, the endonuclease domain deletion mutant IRE1β(Δ783-925) almost as strongly inhibited stress-induced xbp1 splicing, indicating that endonuclease activity is not required either. In cells expressing the double domain deletion mutant IRE1β(Δ453-925), however, though it also inhibited stress-activated IRE1α activity, the extent of inhibition was reduced, and notably less than that observed for the single point substitution kinase-dead mutant IRE1β(K547A). Thus, IRE1β kinase and endonuclease enzymatic activities are not required to suppress stress-induced IRE1α signaling, though the structural domains appear to be needed, suggesting that IRE1β and IRE1α may interact.

To test this idea, we first used the FLAG-tagged IRE1β-inducible HEK293^doxIRE1β^ cell model for co-immunoprecipitation studies. As presented above, when treated with 100 ng/mL Dox, these cells express IRE1β at levels 100-fold above IRE1α (Figure 1D), roughly comparable to the level of IRE1β relative to IRE1α expression observed in mouse primary intestinal goblet cells (Supplemental Figure 1A). Immunoprecipitation of IRE1β from lysates of these cells using anti-FLAG antibody co-immunoprecipitated endogenous IRE1α as assessed by immunoblot (Figure 3D). This was observed only in HEK293^doxIRE1β^ cells expressing IRE1β (there was no detectable co-IP of endogenous IRE1α with anti-FLAG in cells not treated with Dox). Notably, IRE1α and IRE1β co-immunoprecipitated in the absence of thapsigargin (though to a lesser extent), consistent with the possibility that IRE1β may constitutively interact with IRE1α to prevent IRE1α activation. Similar results were obtained using HEK293 cells transiently transfected with FLAG-tagged IRE1β (Supplemental Figure S3A). Thus, IRE1β appears to interact physically with IRE1α.

We next sought biochemical evidence for direct interaction using recombinant proteins in vitro. Full-length myc-tagged IRE1α and IRE1β were expressed and purified from Expi293 cells. Molecular identity of the recombinant proteins was verified by mass spectrometry of Coomassie-stained bands, and by immunoblot using antibodies against IRE1α, IRE1β, or the myc-epitope (Figure 3E). Endonuclease activity was measured in vitro for each of the IRE1 isoforms by incubating the purified enzymes at different concentrations with a fixed amount of a model xbp1 reporter substrate (Wiseman et al., 2010). In this assay, endonuclease cleavage of the xbp1-reporter leads to increased fluorescence, which can be measured in real-time. As reported in the literature for isolated IRE1 cytosolic domains (Imagawa et al., 2008), the endonuclease activities for full-length IRE1α and IRE1β were found to be quite different. Incubation of the reporter substrate with purified full-length IRE1α resulted in a robust increase in fluorescence on the time scale of minutes (Figure 3F, top). Reactions with purified full-length IRE1β, on the other hand, showed no detectable increase in fluorescence when measured on the same time scale (Figure 3F, bottom). The purified IRE1β protein was nonetheless active in cleavage of the RNA substrate, though with much weaker endonuclease activity and slower kinetics as reporter cleavage by IRE1β was eventually detected to much lower extent after hours, not minutes, of incubation (Figure 3F, bottom panel inset). Similar results were obtained in cell-based assays using HEK293/ IRE1α^KO^ cells that do not express IRE1α (Supplemental Figure S3B, top panel) and lack endogenous xbp1 splicing activity (Supplemental Figure S3B, middle and bottom panels). Overexpression of IRE1α in this model resulted in elevated levels of spliced xbp1 transcript that could be significantly enhanced by treatment with thapsigargin or, to a lesser extent, tunicamycin to induce ER stress (Supplemental Figure S3C, purple bars). Overexpression of IRE1β at similar levels induced comparably lower levels of spliced xbp1 transcript that were not enhanced by ER stress stimulation (Supplemental Figure S3C, green bars). Thus, although enzymatically active, IRE1β appears to have weaker endonuclease activity for xbp1 substrates.

When the purified proteins were assayed in vitro under steady state conditions, IRE1α exhibited classical Michaelis-Menten kinetics for cleavage of the xbp1 reporter with reproducibly similar *K*_M_ values from different batches of protein (Prep #1: *K*_M_ = 1.9 ± 0.3 μM; Prep #2: *K*_M_ = 2.0 ± 0.3 μM). When assayed in the presence of IRE1β (at 5-fold molar excess), the maximal reaction velocity was reduced by 15-25% (Figure 3G, light blue curves). The change in *V*_max_ was reproduced in multiple experiments and with two independently prepared batches of purified protein. When analyzed using a noncompetitive inhibitor model, the reduction in *V*_max_ for IRE1α activity caused by IRE1β corresponded to an inhibitory constant of *K*_I_ ≈ 300 nM. Although IRE1β could potentially bind substrate and thus reduce availability of substrate for IRE1α, this would have a negligible effect on the observed IRE1α endonuclease activity. Even assuming stoichiometric binding, the available substrate concentration would only be changed by a few percent for the highest substrate concentrations used and with no effect on the analysis using the noncompetitive inhibitor model. Thus, IRE1β, which has weak endonuclease activity for xbp1, can interact directly with IRE1α to inhibit its endonuclease activity.

### IRE1β has impaired phosphorylation and does not form higher-order oligomers

To a large extent, IRE1β behaved in these studies similarly to a kinase-dead version of IRE1α, which also has weak endonuclease function and has been shown to act as a dominant negative inhibitor of endogenous IRE1α xbp1-splicing (Tirasophon et al., 1998). To gain further insight, we reproduced this experiment. Overexpression of the kinase-dead IRE1α(K599A) mutant in HEK293 cells nearly completely blocked the robust xbp1 splicing reporter activity seen in stress-stimulated control cells with endogenous IRE1α (Figure 4A). This is understood in the literature as a failure of the kinase-dead mutant to *trans* autophosphorylate (Tirasophon et al., 2000) and form higher-ordered IRE1-oliomers required for endonuclease activation (Li et al., 2010). Consistent with those reports, when the IRE1α(K599A) kinase-dead mutant was co-expressed with fluorescently-tagged wild type IRE1α-mCherry in HEK293 cells, the gel filtration elution profile of IRE1α-mCherry was shifted to lower molecular weight fractions with a concomitant loss of in vitro endonuclease activity (Figure 4B). This result phenocopied our results with IRE1β (Figure 2D) and underscores the biologic plausibility that IRE1β, like the kinase-dead IRE1α(K599A) mutant, may act in a dominant negative manner to suppress stress-induced IRE1α signal transduction.

**Figure 4.**
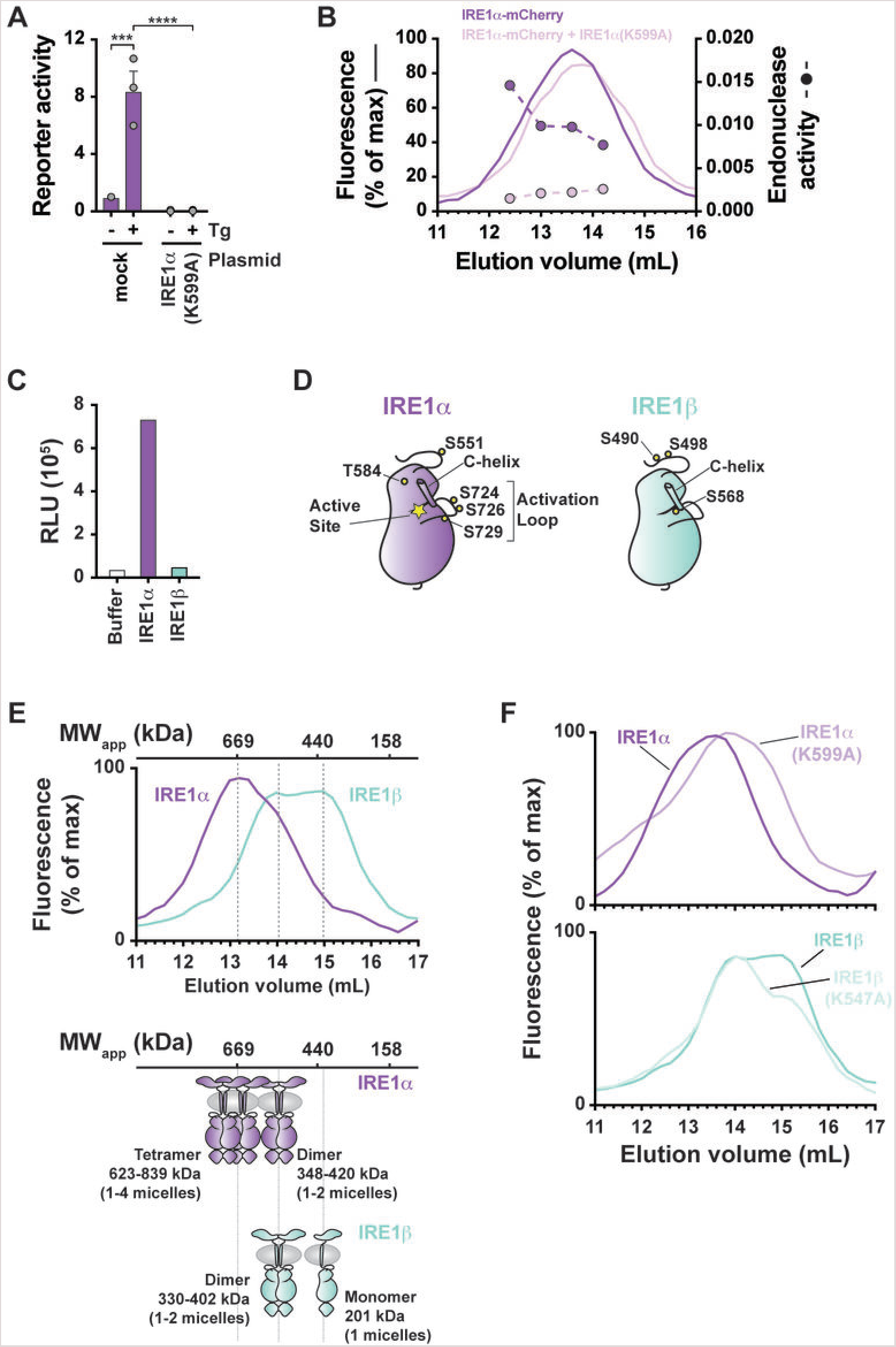
IRE1β has impaired phosphorylation and does not form higher-order oligomers. (A) HEK293 cells were co-transfected with xbp1 splicing reporter and control vector or IRE1α(K599A) expression vector, treated with Tg for 8 hr, and assayed for reporter activity as in Figure 3B. Bars represent mean ± SEM for 3 independent experiments. (B) Same as in Figure 2D for HEK293 cells transfected with IRE1α-mCherry or IRE1α-mCherry + IRE1α(K599A). Cells were lysed in dodecylmaltoside buffer and fractionated by gel filtration chromatography. IRE1α elution was monitored by mCherry fluorescence (solid lines) and endonuclease activity was measured for indicated fractions (symbols and dashed lines). (C) In vitro kinase activity measured for purified full-length IRE1α and IRE1β using ADP-Glo luciferase assay. Bars represent values from a single experiment. Results are representative of two independent experiments that have variable baseline (buffer control) luminescence. (D) Schematic of IRE1α and IRE1β kinase domains illustrates positions of phosphorylation sites detected by mass spectrometry. Indicated sites were detected in two independent preparations of purified protein. (E) HEK293 cells were transfected with IRE1α-mCherry or IRE1β-mCherry expression vectors, lysed in dodecylmaltoside buffer, and fractionated by gel filtration chromatography. IRE1 elution was monitored by mCherry fluorescence. Chromatograms are representative of more than 5 independent experiments. Elution positions of proteins with known molecular weight are indicated above chromatogram. Schematic of putative IRE1 oligomerization states are shown below chromatograms. (F) Same as in (E) for expression and analysis IRE1α-mCherry, IRE1α(K599A)-mCherry, IRE1β-mCherry, and IRE1β(K547A)-mCherry constructs.

The closely similar molecular phenotypes for the kinase-dead IRE1α(K599A) mutant and wild type IRE1β also suggested to us that IRE1β function might be explained by deficient kinase activity and impaired auto-phosphorylation in native IRE1β. To test this idea, we first applied our purified proteins to an autophosphorylation assay in vitro. Kinase activity (autophosphorylation) was measured indirectly by ATP hydrolysis. IRE1β had comparatively much weaker kinase activity than IRE1α (Figure 4C). And, when assessed by mass spectrometry, IRE1α and IRE1β isolated from HEK293 cells had different patterns of phosphorylation. In two independent preparations, IRE1α had phosphorylation on S724, S726, and S729 in the activation loop of the kinase domain – phosphorylation sites that are associated with increased IRE1α endonuclease activity (Prischi et al., 2014). Strikingly, we did not detect any phosphorylation for the equivalent activation loop serine residues in IRE1β (Figure 4D and Supplemental Data S1).

We also found that IRE1β failed to form higher order oligomers when expressed in HEK293 cells. Oligomerization of IRE1β and IRE1α was compared using mCherry-tagged constructs and gel filtration chromatography as described above. When expressed on its own, IRE1α-mCherry eluted from gel filtration as a mix of species with elution peaks around 13.0 mL and 14.0 mL as we found for endogenous IRE1α (Figure 4E, purple tracing). Based on elution of molecular weight standards, we propose these correspond to tetramer and dimer configurations, respectively. In vitro, endonuclease activity predominantly co-eluted with the higher molecular weight tetramer species (Figures 2C and 2D). IRE1β-mCherry on the other hand, when expressed on its own, eluted from gel filtration considerably later than IRE1α at 14 mL and 15 mL, presumably in dimeric and monomeric configurations respectively (Fig 4E, light green tracing). The active higher molecular weight form of IRE1α appeared to be phosphorylation dependent, as the kinase dead IRE1α(K599A) mutant eluted from gel filtration in the lower molecular weight fractions (Figure 4F, top panel), closely similar to IRE1β. And unlike the shift to lower apparent molecular weight seen for the kinase-dead IRE1α mutant, the elution profile for the corresponding kinase-dead IRE1β(K547A) mutant was similar to wild type IRE1β (Figure 4F, bottom panel). Thus, IRE1β appears to have impaired kinase activity and autophosphorylation, at least at sites that are associated with forming higher molecular weight oligomers and that typify the active species for IRE1α.

### A non-conserved amino acid in the kinase domain active site regulates IRE1 endonuclease activity

To identify structural features that might explain the differential kinase, phosphorylation, and oligomerization features of the two IRE1 homologues, we analyzed the primary structure of IRE1α and IRE1β kinase domains. The kinase domains share 80% sequence similarity, including conservation of key catalytic and regulatory residues. However, one divergent site stood out: H692 in the kinase domain active site of human IRE1α is a glycine (G641) in IRE1β (Figure 5A, arrow). In other Ser/Thr kinases, Gln, Glu, or His, typically occupy this position, and contribute side chains that in some instances make contact with the β-phosphate group of bound nucleotide. In the apo, active configuration of IRE1α, the side chain of H692 is oriented toward the flipped-in conformation of the conserved DFG motif (H692 NE2 is 4.8 A from D711 OD2). Thus, this position may be important for either stabilizing an active configuration of IRE1α or enabling nucleotide binding. In either case, any favorable side chain interactions in IRE1α would be lost with a glycine at this position in IRE1β.

**Figure 5.**
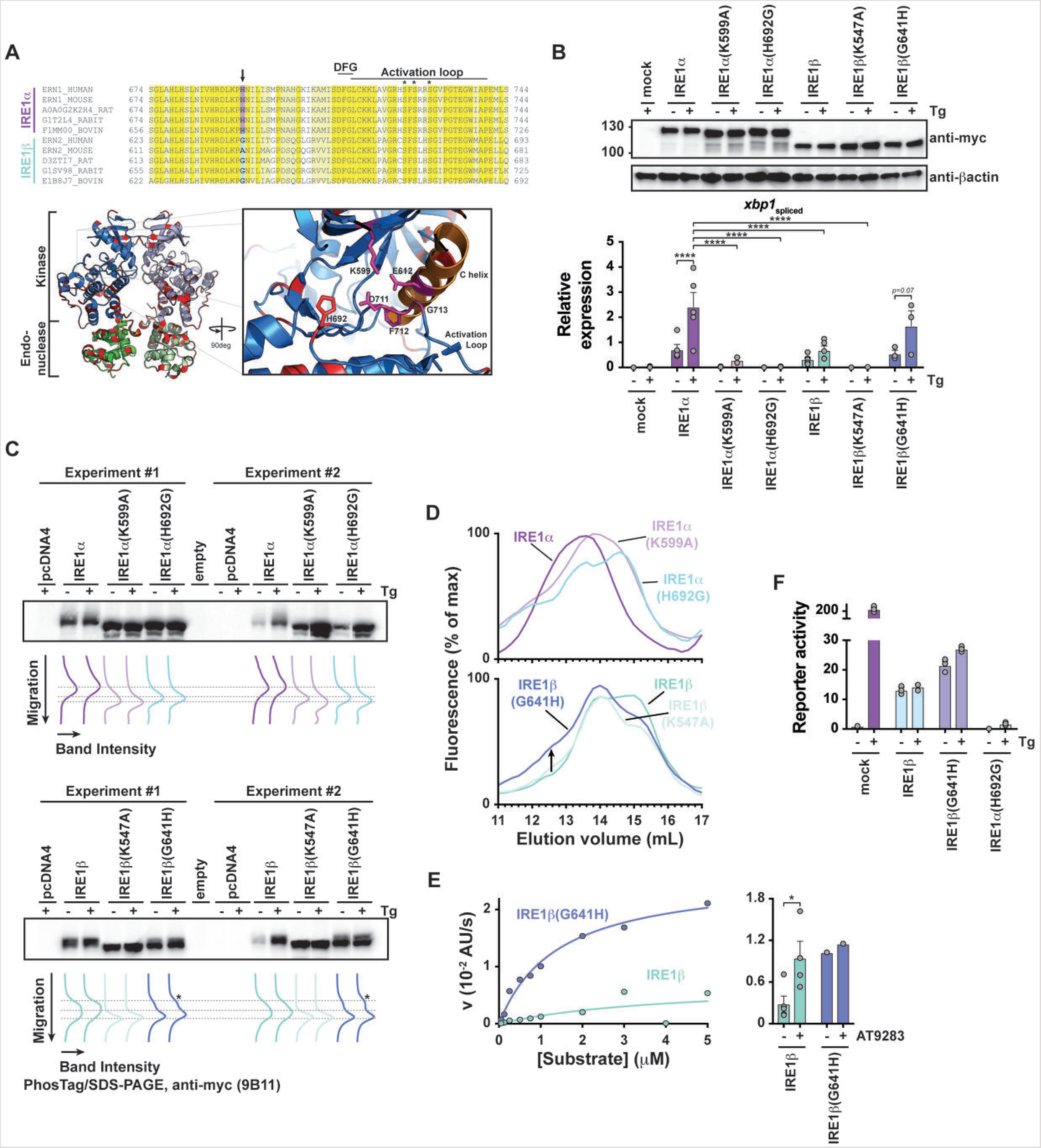
A non-conserved amino acid in the kinase domain active site regulates IRE1 endonuclease activity. (A, top) Sequence alignment for IRE1α and IRE1β active site and activation loop residues. Conserved sequences are colored yellow (identical) and light yellow (similar). The position of non-conserved His→Gly substitution is indicated. (Bottom, left panel) Ribbon diagram of human IRE1α kinase and endonuclease domains (pdb 5hgi, (Feldman et al., 2016)). Residue positions that are not conserved between IRE1α and IRE1β are colored red. (Bottom, right panel) Cartoon illustrates close-up view of kinase domain active site and position of H692 side chain. (B) HEK293/IRE1α^KO^ cells were transfected with indicated IRE1 constructs, treated with thapsigargin (300 nM) for 8 hr, and assayed by western blot for IRE1 expression (anti-myc) or by qPCR for spliced xbp1 transcript. Bars represent mean ± SEM for 3-5 independent experiments. (C) Samples from (B) were assayed by PhosTag/SDS-PAGE and western blot with anti-myc antibody. Band migration position and intensity are plotted under blot. Lines indicate migration position of different phosphorylation species. The blot includes samples from 2 independent experiments. (D) Same as in Figure 4F for indicated IRE1-mCherry constructs. Chromatograms are representative of at least 2 independent experiments. (E, left panel) In vitro endonuclease activity measured for affinity-purified IRE1β and IRE1β(G641H) under steady state conditions. Reaction velocities were measured as a function of substrate concentration using 10 nM enzyme. Lines show best-fit of Michaelis-Menten equation with *K*_M_ = 4 ± 8 μM and *V*_max_ = 0.007 ± 0.008 s^-1^ for IRE1β and *K*_M_ = 1.4 ± 0.2 μM and *V*_max_ = 0.027 ± 0.001 s^-1^ for IRE1β(G641H). (right panel) In vitro endonuclease activity measured for 10 nM enzyme and 1 μM xbp1 reporter substrate in the presence or absence of 100 μM AT9283 kinase inhibitor. Bars represent mean ± SEM for 3 independent experiments. (F) HEK293 cells were co-transfected with xbp1 splicing luciferase reporter and indicated IRE1 constructs, treated with thapsigargin (300 nM) for 4 hr, and assayed for luciferase activity as in Figure 3B. Bars represent mean ± SEM for 3 independent experiments.

**Figure 6.**
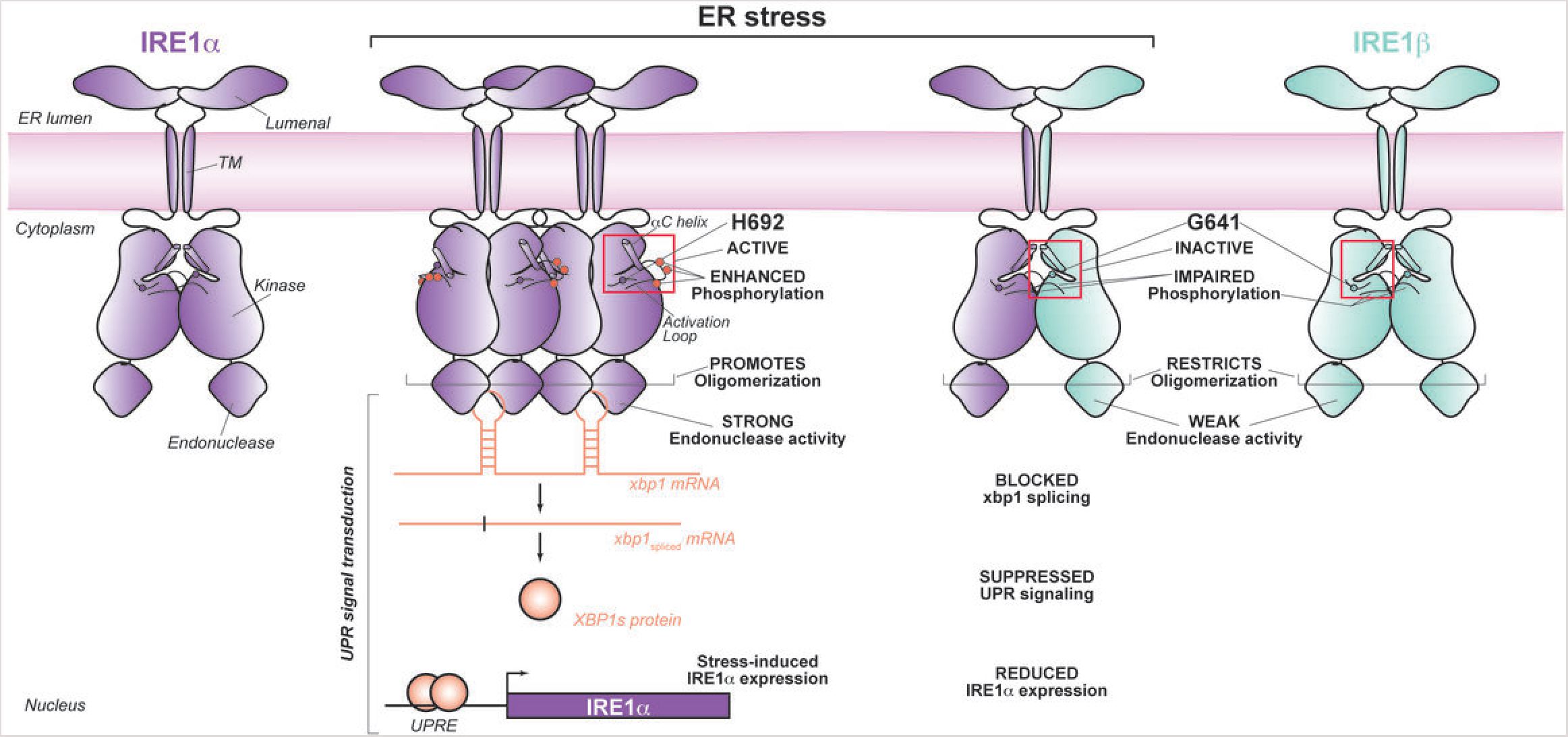
IRE1β functions as a dominant-negative suppressor of IRE1α signaling.

To test if the non-conserved His/Gly in the kinase domain regulates xbp1 splicing activity, we prepared IRE1α(H692G) and IRE1β(G641H) mutant constructs along with kinase-dead versions of each (IRE1α(K599A) and IRE1β(K547A)), expressed them in HEK293^IRE^1^αKO^ cells, and assayed for xbp1 splicing, phosphorylation, and oligomerization. The mutant versions of IRE1α and IRE1β were each expressed at similar levels as their wild type counterparts (Figure 5B, top panel). Strikingly, the H692G mutation in IRE1α completely abolished both basal and stress-induced xbp1 splicing (Figure 5B). This was associated with a faster migrating species on PhosTag SDS-PAGE that was similar to the kinase-dead IRE1α(K599A) mutant (Figure 5C), suggesting that the H692G mutation impaired receptor phosphorylation. Impaired phosphorylation also blocked IRE1α(H692G) oligomerization, as IRE1α(H692G)-mCherry eluted from gel filtration of cell lysates at lower apparent molecular weight, much like the IRE1α(K599A) mutant and wild type IRE1β (Figure 5D).

The corresponding and reversed G641H mutation in IRE1β, on the other hand, rescued stress-stimulated xbp1 splicing (Figure 5B, dark blue bars on far right and Supplemental Figure S5), and IRE1β phosphorylation, as evidence by slower migrating species on PhosTag SDS-PAGE (Figure 5C). The G641H (“rescue”) mutation in IRE1β also partially promoted formation of a higher molecular weight species as assessed by elution upon gel filtration of cell lysates (Figure 5D, bottom panel) and enabled higher levels of IRE1β endonuclease activity *in vitro* (Figure 5E, left panel). Under steady state conditions, IRE1β(G641H) displayed an increased *V*_max_ and reduced *K*_M_, suggesting improved catalytic turnover per unit enzyme and substrate recognition. Interestingly, an IRE1-specific kinase inhibitor that stabilizes an active kinase domain conformation and promotes endonuclease activity (Feldman et al., 2016) was able to stimulate IRE1β endonuclease activity to a similar extent as the G641H mutation (Figure 5E, right panel), suggesting that the His at this position helps stabilize an active kinase domain configuration. Finally, we co-expressed the wild type or IRE1β(G641H) mutant in HEK293 cells containing the xbp-splicing luciferase reporter, and found the IRE1β(G641H) mutant suppressed endogenous IRE1α activity to a lesser extent than wild type IRE1β, although both still had significantly lower reporter activity compared to mock-transfected cells treated with Tg (Figure 5F). The reverse IRE1α(H692G) mutant on the other hand, much like the kinase dead IRE1α(K599A) mutant, completely abolished stress-induced xbp1 splicing by endogenous IRE1α. Together, these data implicate G641 in IRE1β (and H692 in IRE1α) as a critical residue linking the IRE1 kinase domain active site with autophosphorylation, oligomer assembly, xbp1 splicing, and the response to canonical forms of ER stress by dominant-negative interaction with IRE1α.

## DISCUSSION

Our findings delineate a mechanism for down-regulation of the UPR unique to epithelial cells lining mucosal surfaces. The ER of these cells contains IRE1β, which under conventional conditions of ER stress appears to interfere with IRE1α signaling. Upon the induction of ER stress, IRE1β interacts with and inhibits IRE1α in a dominant negative manner. A single non-conserved residue in the kinase domain active site of IRE1β largely explains this activity. Downstream of IRE1α activation, IRE1β also blocks expression of genes regulated by XBP1s, including IRE1α itself. Thus, IRE1α regulates its own expression by producing its own transcription factor. This auto-feedback loop further enables the down-regulatory effects of IRE1β on IRE1α to affect the UPR. These results support the idea that IRE1β has been evolutionarily conserved to dampen the amplified ER stress responses expected of the host-environment interface and required for mucosal host defense.

IRE1β appears to interact directly with IRE1α to block oligomerization and thus activation. Evidence for such a direct interaction includes: (1) the cytosolic domains of IRE1β are structurally required to suppress IRE1α signaling, independently of their kinase and endonuclease activities; (2) IRE1β co-immunoprecipitates with endogenous IRE1α under basal and stress-stimulated conditions; and (3) in biochemical assays using purified proteins and physiologically-relevant stoichiometries, IRE1β non-competitively inhibits the endonuclease activity of IRE1α. Given the overall sequence similarity of IRE1β and IRE1α it is possible that IRE1β could form a heterodimer with IRE1α that mimics an IRE1α homodimer (although this is likely a weaker interaction than between IRE1α(K599A) and IRE1α, which suppresses xbp1 splicing to a greater extent). Further *in vitro* and structural studies are needed to define the molecular interface mediating the IRE1β-IRE1α interaction. Likewise, other ER lumenal and membrane proteins, such as BiP (Bertolotti et al., 2000), ERdj4 (Amin-Wetzel et al., 2017), Hsp47 (Sepulveda et al., 2018), and Sec61 (Sundaram et al., 2017) regulate IRE1α activity, and we have not studied how IRE1β may affect these interactions. And, we do not measure IRE1-dependent RIDD activity (Hollien et al., 2009; Hollien and Weissman, 2006). It is possible this form of signal transduction by IRE1 remains unaffected or even amplified in cells expressing both IRE1 isoforms.

The weak xbp1 splicing activity exhibited by IRE1β enables the protein to act in a dominant negative manner blocking IRE1α xbp1 splicing. This is largely explained by a non-conserved mutation in the IRE1β kinase domain active site that links impaired autophosphorylation and oligomerization with inhibition of IRE1α. Exactly how the G641 residue regulates IRE1β kinase activity and ultimately endonuclease outputs however remains inconclusively explained. Since the small molecule kinase inhibitor AT9283 acts to rescue the endonuclease activity of IRE1β, presumably by stabilizing the active conformation of the kinase domain as it does in IRE1α (Feldman et al., 2016), we propose the G641 residue acts to disable the conformational stability of the kinase domain. As a general rule, kinase domains are well known to exhibit defined transitions between active and inactive configurations regulated by phosphorylation, nucleotide and substrate binding, and higher-order protein dimerization/oligomerization. In some receptor kinases, the conformation of the kinase domain active site has been linked to receptor assembly and signal transduction (Mi et al., 2011). This appears to be the case for both IRE1 isoforms. In the case of IRE1β, the destabilized kinase domain active conformation, we propose, accounts for both its weak xbp1 splicing activity and its apparent lack of response to activators of ER stress. Rescue of the G641 residue by substitution with His, however, does not completely revert IRE1β to full IRE1α-like activity. There are other notable structural differences between the two proteins, such as in the immediate juxtamembrane and likely lumenal domain regions that also may affect protein function in physiologically relevant ways.

As a consequence of inhibiting IRE1α endonuclease activity, IRE1β attenuates the production of the well-described transcription factor XBP1s. Here, we find that XBP1s also regulates IRE1α transcription, thus elucidating how IRE1α is induced by the UPR. Since XBP1s binds promoters of genes throughout the genome, the effect of IRE1β on basal and stress-induced xbp1 splicing will likely have an impact on many gene expression programs. With respect to the UPR, genome-wide chromatin immunoprecipitation (ChIP) analysis of muscle and secretory cells showed that XBP1s binds promoter regions for *hspa5* (BiP), *atf4*, and *ddit3* (CHOP), and other genes responsive to ER stress. Thus, it is possible that by inhibiting IRE1α endonuclease activity, IRE1β could reprogram the transcriptional output of XBP1s and potentially affect all arms of the UPR.

We also propose that a function of IRE1β to suppress stress-induced IRE1α signaling may be fundamentally important for maintenance of intestinal physiology. The surface of epithelial cells facing the intestinal lumen is exposed to a diverse array of environmental factors that may cause cell damage—such as some dietary components, orally-administered drugs, bacterial toxins, and pathogenic and commensal microorganisms or their products. Each of these could potentially activate or modify cellular stress responses, directly or indirectly to affect the UPR. As unresolved ER stress and prolonged UPR signaling induces cell death, such responses left unchecked are likely harmful to the mucosal barrier epithelium. Our results explain the mechanism of action and concur with the idea that IRE1β has been conserved to buffer IRE1α and XBP1s signal transduction against stimuli encountered by epithelial cells at the host-environment interface.

## MATERIALS AND METHODS

### Caco2 and T84 cell culture

Caco2BBE cells were maintained in DMEM supplemented with 15% FBS and T84 cells were maintained in 1:1 DMEM/F12 media supplemented with 6% newborn calf serum. Cells were plated on 1 cm^2^ Transwell inserts (0.4 um pore size polycarbonate membranes for Caco2 cells and 3 um pore size polyester membranes for T84 cells) and allowed to polarize for 7 days. Trans-epithelial electrical resistance was measured using Epithelial Volt/Ohm Meter (EVOM, World Precision Instruments) to assess monolayer formation. Monolayers were treated with thapsigargin (3 μM final) or DMSO as control in apical and basolateral compartments for 2 hr at 37 °C. Cells were washed 2X with ice-cold PBS and stored at −80 °C.

### Colonoid culture

All housing and procedures involving live vertebrate animals were reviewed and approved by Boston Children’s Hospital Institutional Animal Care and Use Committee. IRE1β^+/+^ and IRE1β^-/-^ mice were euthanized. The colon was excised, fecal pellets were removed, and the tissue was flushed with 5 mL of ice-cold PBS. The colon was opened longitudinally and cut into several small (2-3 mm) pieces. Tissue pieces were washed 2-3 times in PBS and placed in PBS with 10 mM EDTA for 45 min with end-over-end rotation at 4 °C. Crypts were dissociated by vigorous shaking for 5-7 min, and the supernatant was collected and diluted 2-fold with base media (Advanced DMEM/F12 supplemented with 20% FBS, 10 mM HEPES, 1X Glutamax, and 1X Pen/Strep). Tissue pieces were incubated a second time with PBS/EDTA and dissociated cells collected in the same manner. Collected cells were passed through a 100 μM strainer followed by a 40 μM strainer. Cells retained on the 40 μM strainer were washed with base media, collected with 10 mL base media, and spun down at 300xg for 3 min. Crypts were resuspended in Matrigel (on ice) and 30 μL drops were plated in 24-well plates. Plates were inverted and incubated at 37 °C to polymerize Matrigel. Complete media (Base media + 50% WRN-conditioned media (Miyoshi and Stappenbeck, 2013)) was added to each well and cultures were incubated at 37 °C 5% CO2. Media was changed every other day. Cultures were passaged every 4-7 days as needed. Matrigel was dissolved using Cell Recovery Solution (Corning), cells were enzymatically dissociated with trypsin, and the cell pellet was resuspended in 1.5-2-fold more Matrigel than used in the previous plating. For experiments, colonoids were plated in triplicate. At 5 days post-plating, colonoids were treated with SubA_A272_B or SubAB (100 ng/mL) for 24 hr. Matrigel drops were collected and pooled, colonoids recovered using Cell Recovery Solution, washed 1X with PBS, and stored at −80 °C.

### HEK293^doxIRE1β^ cell model

HEK293^doxIRE1β^ cells were established by subsequent transduction of low passage HEK293 cells (obtained from ECACC) with lentiviral particles encoding the pLenti3.3 doxycycline-controlled regulator and pLenti6.3/V5/ERN2-FLAG (Invitrogen). ERN2-FLAG coding sequence was introduced by Gateway cloning. All sequences were confirmed by sequencing. Transduction was performed with concentrated and tittered virus at multiplicity of infection (MOI) of 1 for pLenti3.3 and MOI of 2 for pLenti6.3, respectively.

### Overexpression of IRE1 constructs by transient transfection

General molecular biology reagents, including restriction enzymes, Phusion HF Mastermix, T4 PNK, CIP, Quick Ligation Kit, and DH5α competent cells were from New England Biolabs. GeneJet DNA purification kits were from ThermoFisher. pcDNA4-MycHisB plasmid (ThermoFisher) was digested with *HindIII* and *EcoRI*. Oligonucleotides with *HindIII* restriction sites-Kozak sequence-start codon-murine Igκ leader sequence-FLAG tag-*EcoRI* restriction site were annealed, phosphorylated with T4 PNK, and ligated into *HindIII/EcoRI*-digested pcDNA4-MycHisB vector to create pSecFLAG-MycHis for in-frame cloning with C-terminal myc epitope and His-tag. Additional pSecFLAG vectors were created with mCherry coding sequence inserted between NotI and XbaI restriction sites of pSecFLAG (for in-frame cloning with C-terminal mCherry tag, pSecFLAG-mCherry) or a streptavidin binding peptide (SBP) tag immediately upstream of myc epitope (pSecFLAG-SBP-MycHis). IRE1α (S24-L977), IRE1β (L35-R925), and IRE1β deletion construct (L35-S782, L35-R452) coding sequences were PCR amplified from IRE1α and IRE1β templates (provided by K. Khono) using primers with *EcoRI* and *NotI* restriction sites. PCR-amplified products were digested and ligated into *EcoRI/NotI*-digested pSecFLAG-MycHis, pSecFLAG-mCherry, or pSecFLAG-SBP-MycHis vectors. IRE1α(K599A), IRE1α(H692G), IRE1β(K547A), and IRE1β(G641H) mutations were introduced using QuickChange Lightning Site-Directed Mutagenesis Kit (Agilent). All constructs were confirmed by restriction digest and sequencing (Harvard Biopolymer Facility).

HEK293T, HEK293, and HEK293IRE1α^KO^ cells were maintained in DMEM supplemented with 10% FBS. Cells were seeded in 24-well plates at a density of 2 x 10^5^ cells/well in 0.5 mL media. At 24 hr after plating, cells were transfected with IRE1 expression plasmids or pcDNA4 as control (500 ng per well) using PEI (linear 25 kDa, Polysciences, Inc.) at a DNA:PEI mass ratio of 1:3. At 18-24 hr post-transfection, cells were treated with thapsigargin or SubAB along with appropriate controls as described in figure legends. After treatment, cells were washed 2X with ice-cold PBS and used for RNA extraction or preparation of whole cell lysates.

### Expression analysis by qPCR

Total RNA was extracted from cell lines and colonoids using the RNeasy Mini Kit (Qiagen). Cell pellets were lysed in Buffer RLT, homogenized with QiaShredder (Qiagen), and processed according to manufacturer’s protocol (including on-column DNase digest). Total RNA concentrations were measured by absorbance at 260 nm and quality was assessed by A260/A280 ratios. Total RNA (typically 500 ng) was used as template for cDNA synthesis using iScript cDNA synthesis kit (BioRad). Target transcripts were amplified using primers listed in Table 1 and Sso Advanced Universal SYBR Green Supermix according to manufacturer’s protocol (BioRad). All qPCR reactions were assayed in triplicate for each sample and the average Cq value was used to calculate the mean expression ratio of the test sample compared to the control sample (i.e. stress-treated compared to control-treated) using the 2-ΔΔCt method. Cq values for targets were analyzed relative to Cq values for *hprt, ppia*, and *gapdh* housekeeping genes.

**Table 1.**
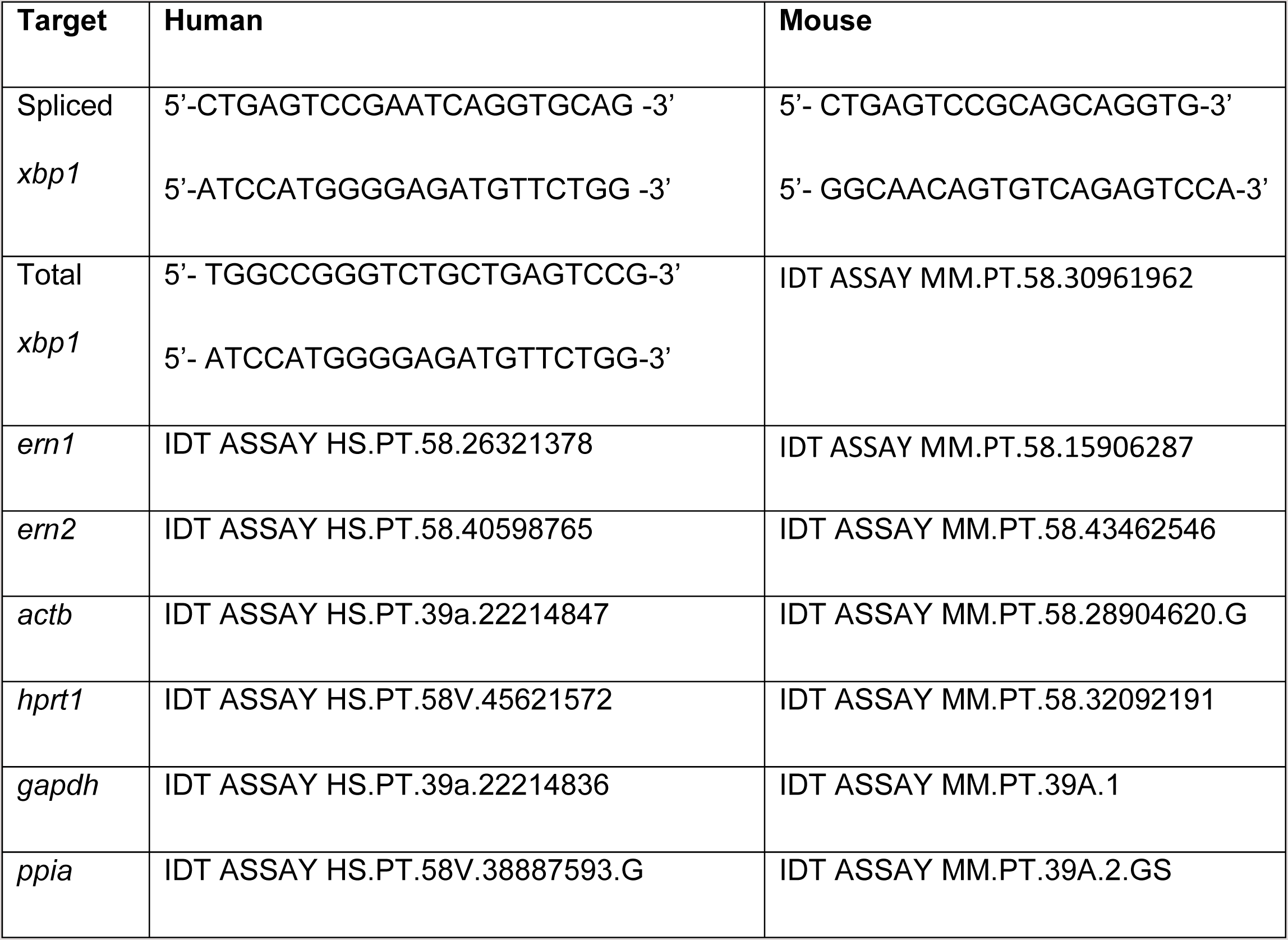
Primers used for qPCR expression analysis.

### Western blots

Whole cell lysates were prepared in RIPA buffer supplemented with Complete Protease Inhibitor (100 μL per cm^2^ surface area of cells). Cells were lysed on ice for 15 min. Lysates were cleared by centrifugation at 16,800xg for 10 min. Total protein was measuring using BCA assay. Typically 10-25 μg total protein was separated on reducing SDS-PAGE (4-15% or 4-20% gradient gels, BioRad). Proteins were transferred to nitrocellulose membranes, blocked with 5% dry milk in TBS/T buffer. Membranes were incubated with primary antibody (diluted in 5% milk, TBS/T) for 1 hr at room temperature or 16-24 hr at 4 °C, washed 3X with TBS/T, and incubated HRP-conjugated secondary antibody (diluted in 5% milk, TBS/T) for 1 hr at room temperature. Membranes were washed 3X with TBS/T, developed with SuperSignal West Femto Maximum Sensitivity Substrate (ThermoFisher), and imaged using Azure c300 Chemiluminescent Western Blot Imaging System (Azure Biosystems). Membranes were stripped with Restore Stripping Buffer (ThermoFisher) and re-probed as needed. Band intensities were quantified using ImageJ. Antibodies used include anti-IRE1α (14C10, Cell Signaling Technologies, 1:1000), anti-IRE1β (PA5-13921, Thermo Fisher, 1:2000), anti-myc tag (9B11, Cell Signaling Technologies, 1:2000), anti-FLAG (M2 F1804, Sigma, 1:1000), anti-XBP1 (M186, sc-7160 Santa Cruz Biotechnology, 1:200), and anti-βactin (AC-15, A5441 Sigma, 1:5000). Secondary antibodies used include HRP-conjugated anti-rabbit IgG (A6154, Sigma, 1:5000 – 1:10000) and HRP-conjugated anti-mouse IgG (A4416, Sigma, 1:5000-1:10000).

### Luciferase reporter assays

pCAX-HA-2xXBP1-Luc-F (xbp1 splicing reporter) was provided by T. Iwawaki (Iwawaki and Akai, 2006). pGL3-5XUPRE-Luc was provided by Ron Prywes (Addgene plasmid #11976, (Wang et al., 2000)). hIRE1α_-1153/+114_-Luc and hIRE1α_-_ 300/+114-Luc reporter vectors were prepared by PCR amplifying the specified IRE1α promoter region from a synthetic gBlock Gene Fragment (IDT) with *NheI* and *NcoI* restriction sites. Restriction-digested, gel purified-PCR products were ligated into *NheI/NcoI*-digested pGL3-basic vector (Promega) immediately upstream of the luciferase start codon. HEK293 cells (0.5 x 10^5^ cells/well) were seeded in 96-well plates and reverse transfected with protein expression vectors and inducible luciferase reporter vectors at a 1:1 ratio with 100 ng total DNA per well. At 18-24 hr post-transfection, cells were treated for indicated times, lysed, and assayed for luciferase activity using Bright-Glo luciferase assay system (Promega). Luminescence was measured using Tecan Spark 10M plate reader. The background signal from untransfected cells was subtracted from all data.

### Co-immunoprecipitation studies

For the inducible cell line, 8 x 10^6^ HEK293^doxIRE1β^ cells were seeded in 100 mm dishes. The next day, IRE1β-FLAG expression was induced with 100 ng/mL DOX for 24 hr. For transient overexpression, 8.2 x 10^6^ HEK293T cells were seeded in 100 mm dishes. The next day, IRE1β-FLAG plasmid was transfected using PEI at a 1:5 DNA:PEI ratio (5 μg DNA). Cells were collected after 24 hr. Cells were lysed in lysis buffer (0.1% NP40, 10% glycerol, 250 mM NaCl, 20 mM HEPES pH 7.9, 1 mM EDTA) with protease and phosphatase inhibitors. Samples were incubated with 10 μL pre-washed Dynabeads T1 (Invitrogen) that were either pre-bound with BioM2 anti-FLAG (Sigma, 1 μL antibody/10 μL beads) or no antibody for 1 hr at 4 °C. After washing with lysis buffer, bound proteins were eluted directly in sample buffer and samples were separated on a 4-12% Criterion Bis-Tris gel (BioRad). Proteins were transferred to PVDF membrane (Immobilon-FL, Millipore), and incubate with anti-IRE1a (1/1000, 14C10, Cell Signaling Technology), M2 anti-FLAG (1/1000, Sigma), and anti-tubulin (1/2000, Sigma). Proteins were revealed with anti-mouse and anti-rabbit HRP conjugated antibodies (1/15000, Jackson Immuno AffiniPure).

### IRE1 oligomerization assays

To assay oligomerization of endogenous IRE1α, cells were resuspended in 25 mM Tris pH 8.0, 150 mM NaCl, 20 mM dodecylmaltoside, 5 mM β-mercaptoethanol, 1X Complete Protease Inhibitor, and 1X PhosSTOP (0.5 mL lysis buffer per 10 cm^2^ surface area of cells), transferred to 1.5 mL tube, and lysed at 4 °C with end-over-end rotation for 1 hr. Lysate was cleared by centrifugation at 16,000xg for 10 min. 250 μL of cleared lysate was fractionated on Superose6 Increase 10/30 gel filtration column (GE Healthcare) equilibrated in running buffer (25 mM Tris pH 8.0, 150 mM NaCl, 0.5 mM dodecylmaltoside, 5 mM β-mercaptoethanol) at 0.5 mL/min. Fractions (0.2 mL) were collected in a 96-well plate from 6 mL to 24 mL elution. Fractions were assayed by SDS-PAGE and western blot with anti-IRE1α antibody to detect endogenous IRE1α.

HEK293 cells were plated at 8 x 10^5^ cells/well in 6-well plates. The next day, pSecFLAG-IRE1α-mCherry or pSecFLAG-IRE1β-mCherry constructs were expressed by transient transfection using PEI at a 1:3 DNA:PEI ratio (2.5 μg total DNA per well). In cases where IRE1-mCherry constructs were co-transfected with unlabeled IRE1 constructs, they were included at equal amounts or co-transfected with pcDNA4 as a control. At 24 hr post-transfection, cells were collected, lysed, and fractionated as described above. Fractions were assayed by measuring mCherry fluorescence using Tecan Spark 10M plate reader (excitation at 570 nm and emission at 620 nm) to detect elution of IRE1-mCherry. Background fluorescence was subtracted, if needed, by taking the average of the first 10 fractions and last 10 fractions. For reinjection experiments (Figure 2C), defined fractions (HMW or LMW) from the original fractionated lysate were re-injected on Superose6 column. IRE1-mCherry elution was monitored by fluorescence. Endonuclease activity was monitored for each fraction in 96-well format as described below using 1 μM substrate and measuring fluorescence at 60 min time point.

### Expression and purification of full-length IRE1 constructs

Full length IRE1 constructs for purification were expressed in the Expi293 Expression System (ThermoFisher). Expi293F cells were maintained and expanded in Expi293 Expression Medium. Cells were transfected with pSecFLAG-IRE1-SBP-MycHis expression plasmids according to the manufacturer’s protocol using 1 μg DNA per mL of culture volume. At 48 hr post-transfection, cells were collected, washed 1X with ice-cold PBS, and resuspended at ∼20 x 10^6^ cells/mL in lysis buffer (25 mM Tris pH 8.0, 150 mM NaCl, 20 mM dodecylmaltoside, 5 mM β-mercaptoethanol, 1x Complete Protease Inhibitor (Sigma) and 1x PhosSTOP phosphatase inhibitor (Sigma)). Cells were lysed for 1 hr at 4 °C with end-over-end rotation. Lysate was cleared by centrifugation at 30,000xg for 15 min. Cleared lysate was applied by gravity flow to a 0.5 – 1.0 mL StrepTactin Sepharose column equilibrated in wash buffer (25 mM Tris pH 8.0, 300 mM NaCl, 1 mM dodecylmaltoside, 5 mM β-mercaptoethanol). Column was washed 4x with 5 mL of wash buffer. Bound protein was eluted with 8x 0.5-column volumes of elution buffer (25 mM Tris pH 8.0, 150 mM NaCl, 1 mM dodecylmaltoside, 5 mM β-mercaptoethanol, 2.5 mM desthiobiotin). Samples of elution fractions were assayed by SDS-PAGE and detected by colloidal Coomassie staining, or by western blot with anti-IRE1α, anti-IRE1β, or anti-myc antibodies. Coomassie stained bands were excised, digested with trypsin, and assayed by LC/MS/MS for protein identification and phosphopeptide mapping (Taplin Mass Spectrometry Facility, Harvard Medical School). In some cases further purification of samples was performed by gel filtration chromatography on Superose 6 column equilibrated in 25 mM Tris pH 8.0, 150 mM NaCl, 0.5 mM dodecylmaltoside, 5 mM β-mercaptoethanol.

### In vitro endonuclease assays

An xbp1 RNA stem loop model substrate was synthesized with a 5’-FAM and 3’-BlackHoleQuencher (IDT): 5’-FAM-CAUGUCCGCAGCGCAUG-BHG-3’ (Wiseman et al., 2010). Endonuclease assays were performed in 96-well plates (80-100 μL reaction volume) or 384-well plates (20 μL reaction volume). Purified enzyme or gel filtration fractions were diluted 2-fold with 2x endonuclease reaction buffer containing substrate (at indicated concentrations) to give final conditions of 50 mM Tris pH 7.5, 150 mM NaCl, 10 mM MgCl_2_, 1 mM ATP, 10 mM DTT, 0.5 mM dodecylmaltoside, and 1X substrate. Formation of cleaved reaction product was monitored by fluorescence using Perkin Elmer VictorX or Tecan Spark 10M plate reader with excitation at 485 nm and emission at 535 nm. In all cases, background was corrected using a buffer-only (no enzyme) control reaction. For steady state kinetic measurements, reaction velocities were determined from the initial change in fluorescence as a function of time and velocity versus [Substrate] data were analyzed using Michaelis-Menten or enzyme inhibitor models in Prism.

### In vitro kinase assays

Purified IRE1 enzyme was incubated with 2x kinase buffer in a final volume of 25 μL (125 nM enzyme, 70 mM Tris pH 7.6, 75 mM NaCl, 10 mM MgCl_2_, 5 mM DTT, 1 mM ATP, 0.5 mM dodecylmaltoside) for 30 min at room temperature. ADP produced from autophosphorylation was detected using the ADP-Glo Kinase Assay (Promega). Luminescence intensity was measured on a Perkin Elmer VictorX plate reader. A no-enzyme control (buffer only) was used to control for the depletion of ATP in kinase reaction prior to detection of ADP.

### Statistical Analysis

In most cases, data consisted of 3 or more independent experiments. Unless otherwise indicated in figure legend, figures include all independent measures shown as symbols and bars represent mean values ± SEM. Mean values were compared using one-way ANOVA for 3 or more groups with multiple comparisons corrected using statistical hypothesis testing (Sidak). Mean values for 2 groups were compared using unpaired student t-test. In figures, significance is indicated by * (p < 0.05), ** (p < 0.01), *** (p < 0.001), **** (p < 0.0001). All analysis were performed in Prism (GraphPad Software).

## ACKNOWLEDGEMENTS

We thank members of the Lencer, Eyckerman, and Janssens labs for critical evaluation and discussion throughout the course of this project. We thank Dr. Kenji Kohno for providing pCAG-hIRE1β plasmid; Takao Iwawaki for providing pCAX-HA-2xXBP1ΔDBD(anATG)-LUC-F xbp1 splicing reporter plasmid; Ron Prywes for providing p5xATF6-GL3 (UPRE-LUC, Addgene plasmid #11976); Qingbo Xu and Lingfang Zeng for providing pCMV5-FLAG-XBP1s plasmid (Addgene plasmid #63680); Dan Chinnapen and Elisha Fielding for providing Expi293F cells and expression protocols; and Dustin Maly for providing AT9283 IRE1 kinase inhibitor and valuable discussions on IRE1 kinase domains and activation. This work was supported by National Institutes of Health grants DK048106, DK084424, and the Harvard Digestive Disease Center P30DK034854 (Core C) (WIL).

## AUTHOR CONTRIBUTIONS

MJG and WIL conceived the project and MJG, WIL, EC, SE, and SJ designed experiments. MJG, EC, and MSS carried out all experiments with technical support from NL, HD, and DDS. YVS, AWP, and JCP provided critical reagents. PL, JRT, and MAS provided critical reagents and intellectual support in experiment design and analysis. MJG, EC, MS, SE, SJ, and WIL analyzed and provided interpretation for all data. MJG and WIL wrote the manuscript with substantial input from EC, MSS, SE, and SJ. Final manuscript and data was made available to all authors for review and comment prior to submission.

## SUPPLEMENTAL FIGURE LEGENDS

**Figure S1 (Supports Figure 1).**
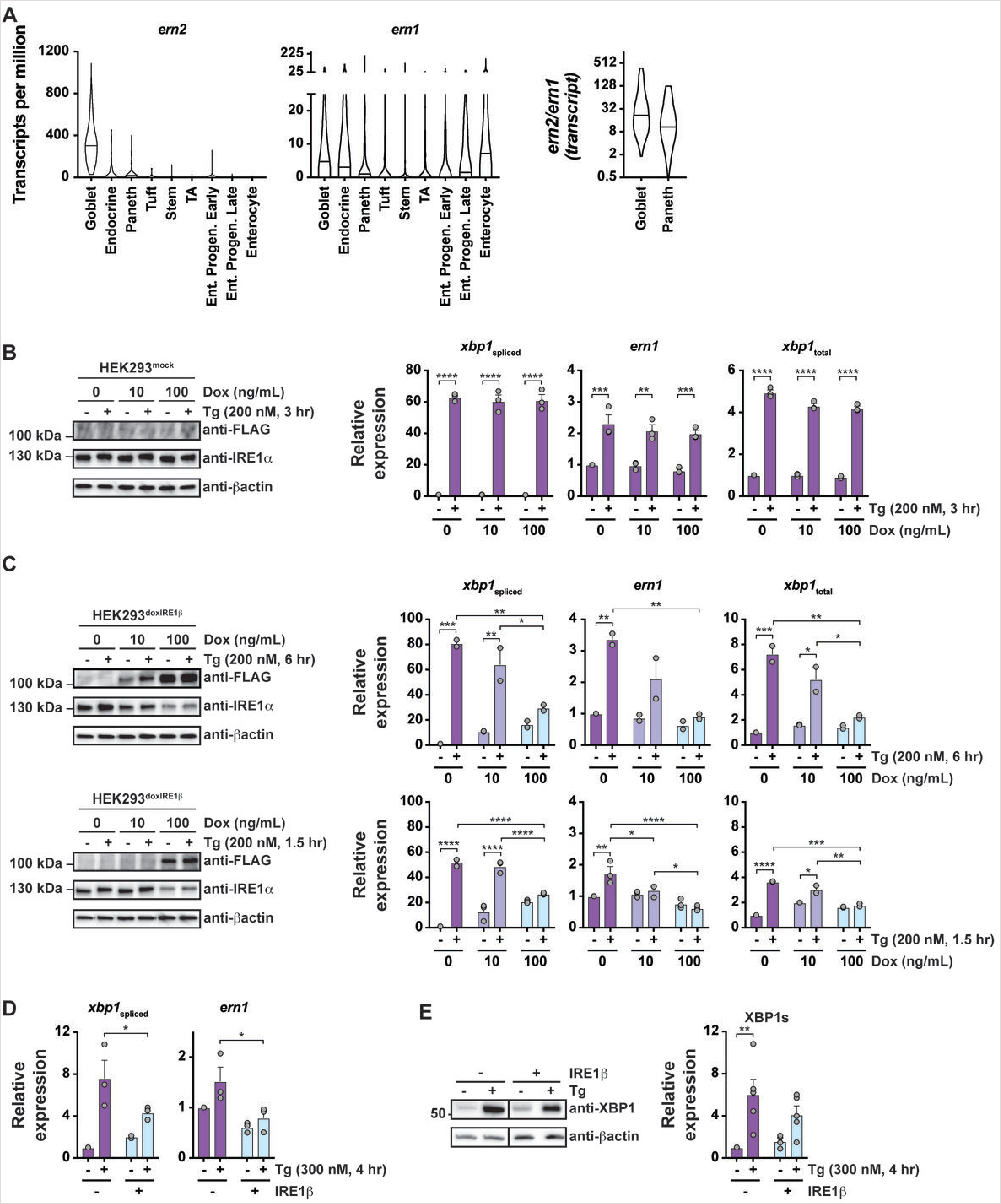
(A) Violin plots for expression of *ern2* and *ern1* transcripts in transcriptionally defined cell populations from mouse small intestine. Lines indicated median values. Data from GSE92332 (Haber et al., 2017). The ratio of *ern2*/*ern1* expression for individual cells is plotted for goblet and Paneth cell populations. (B) HEK293^mock^ cell line was treated with indicated concentration of doxycycline (Dox) for 24 hr. Cells were stimulated with 200 nM thapsigargin for 3 hr and assayed for (left) IRE1β-FLAG and IRE1α protein expression by western blot or (right) *ern1*, total *xbp1*, and spliced *xbp1* transcripts by qPCR. The blot is representative of two independent experiments, and the bars represent mean ± SEM for 3 independent experiments. (C) IRE1β expression was induced in HEK293^doxIRE1β^ cell line with indicated concentration of doxycycline (Dox) for 24 hr. Cells were stimulated with 200 nM thapsigargin for (top) 6 hr or (bottom) 1.5 hr and assayed for (left) IRE1β-FLAG and IRE1α protein expression by western blot or (right) *ern1*, total *xbp1*, and spliced *xbp1* transcripts by qPCR. The blots are representative of 3 independent experiments. The bars represent mean ± SEM for 2 (6 hr) or 3 (1.5 hr) independent experiments. (D) HEK293 cells were transiently transfected with control plasmid or IRE1β expression plasmid, treated with thapsigargin for 4 hr, and assayed for *ern1* and spliced xbp1 transcript by qPCR. Bars represent mean ± SEM for 3 independent experiments. (D) HEK293 cells were transfected with control vector or IRE1β expression vector, treated with thapsigargin for 4 hr, and assayed by western blot for XBP1s protein. A representative blot is shown for anti-XBP1 and anti-βactin antibodies. Bars represent mean ± SEM for normalized band intensity (relative to β-actin) for 5 independent experiments.

**Figure S3 (Supports Figure 3).**
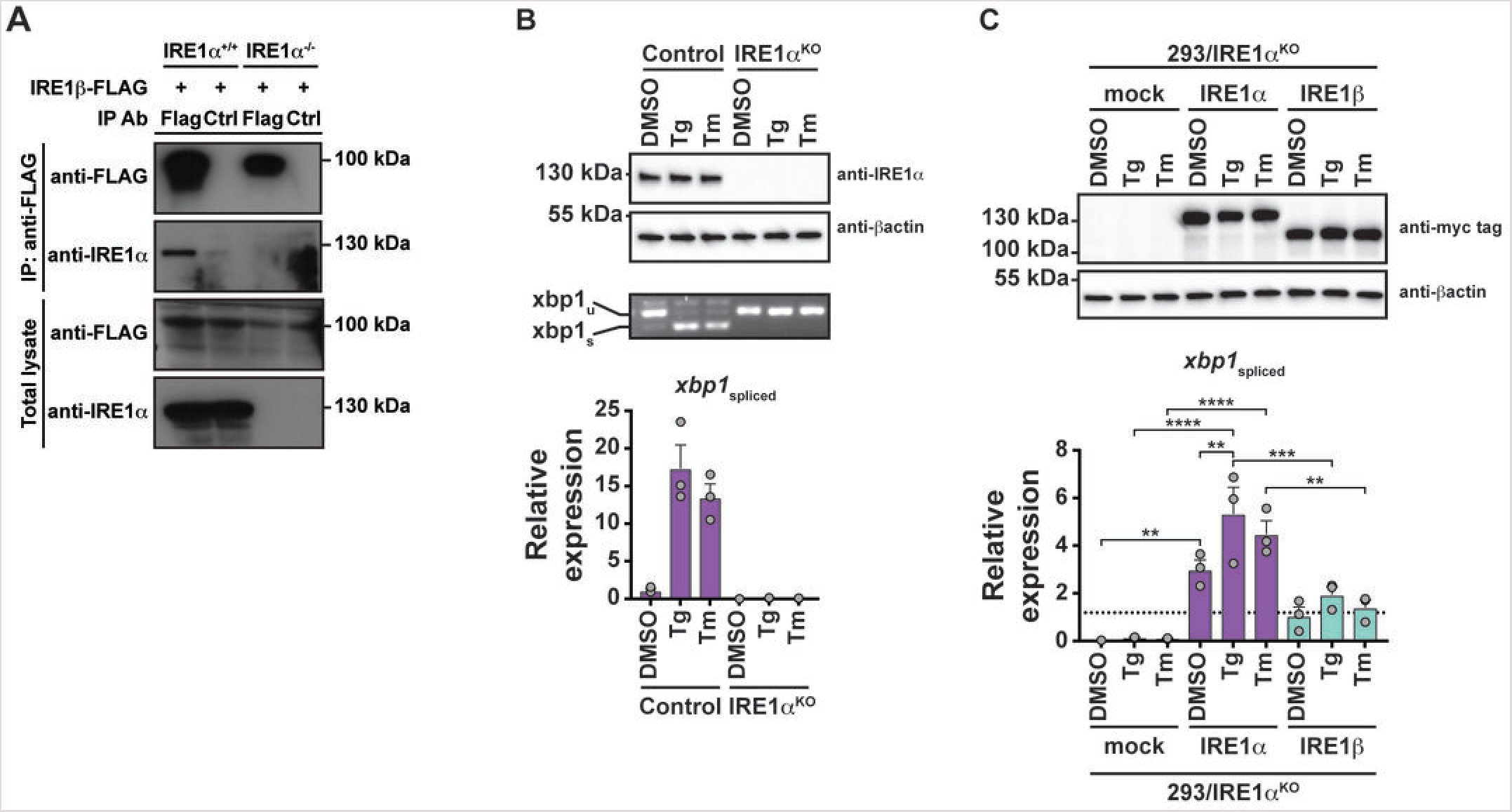
(A) Western blots of anti-FLAG immunoprecipitated samples or total lysates from HEK293 or HEK293/IRE1α^KO^ cells transfected with IRE1β-FLAG expression construct. Endogenous IRE1α was detected with anti-IRE1α antibody and IRE1β was detected with anti-FLAG antibody. (B) HEK293 or HEK293/IRE1α^KO^ cells were treated with DMSO, thapsigargin, or tunicamycin and assayed by (top) western blot for IRE1a expression, (middle) PCR for spliced and unspliced xbp1 transcript, and (bottom) qPCR for spliced xbp1 transcript. Bars represent mean ± SEM for 3 independent experiments. (C) HEK293/IRE1α^KO^ cells were transfected with control vector or IRE1 expression vectors, treated with DMSO, thapsigargin, or tunicamycin, and assayed by (top) western blot for IRE1 expression with anti-myc antibody and (bottom) qPCR for spliced xbp1 transcript. Bars represent mean ± SEM for 3 independent experiments.

**Figure S5 (Supports Figure 5b).**
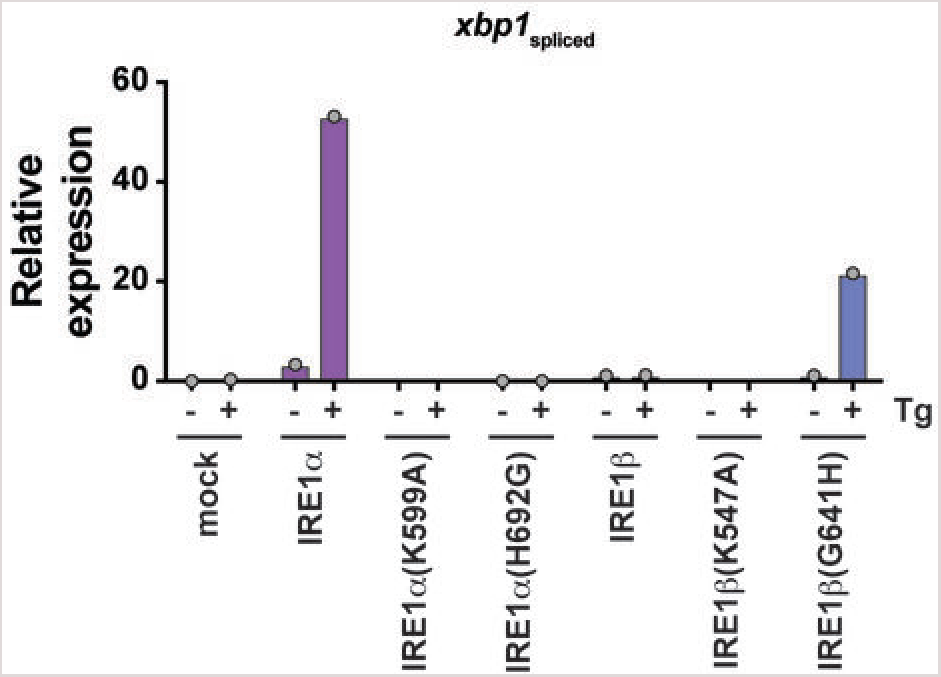
HEK293/IRE1α^KO^ cells were transfected with indicated IRE1 constructs, treated with thapsigargin (300 nM) for 8 hr, and assayed by qPCR for spliced xbp1 transcript. Bars represent a single independent experiment. These data were not included in Figure 5b because the response measured for IRE1α- and IRE1β(G641H)-transfected cells in this experiment was about 10-fold greater in magnitude than any of the other experiments. Although not included in the analysis in Figure 5b, the results from this experiment are still consistent with the conclusion that IRE1α(H692G) abolishes xbp1 splicing and IRE1β(G641H) rescues stress-induced xbp1 splicing activity.

## REFERENCES

Amin-Wetzel, N., Saunders, R.A., Kamphuis, M.J., Rato, C., Preissler, S., Harding, H.P., and Ron, D. (2017). A J-Protein Co-chaperone Recruits BiP to Monomerize IRE1 and Repress the Unfolded Protein Response. Cell 171, 1625-1637.e1613.

Bertolotti, A., Wang, X., Novoa, I., Jungreis, R., Schlessinger, K., Cho, J.H., West, A.B., and Ron, D. (2001). Increased sensitivity to dextran sodium sulfate colitis in IRE1beta-deficient mice. The Journal of clinical investigation 107, 585–593.

Bertolotti, A., Zhang, Y., Hendershot, L.M., Harding, H.P., and Ron, D. (2000). Dynamic interaction of BiP and ER stress transducers in the unfolded-protein response. Nature cell biology 2, 326–332.

Calfon, M., Zeng, H., Urano, F., Till, J.H., Hubbard, S.R., Harding, H.P., Clark, S.G., and Ron, D. (2002). IRE1 couples endoplasmic reticulum load to secretory capacity by processing the XBP-1 mRNA. Nature 415, 92–96.

Feldman, H.C., Tong, M., Wang, L., Meza-Acevedo, R., Gobillot, T.A., Lebedev, I., Gliedt, M.J., Hari, S.B., Mitra, A.K., Backes, B.J., et al. (2016). Structural and Functional Analysis of the Allosteric Inhibition of IRE1alpha with ATP-Competitive Ligands. ACS chemical biology 11, 2195–2205.

Haber, A.L., Biton, M., Rogel, N., Herbst, R.H., Shekhar, K., Smillie, C., Burgin, G., Delorey, T.M., Howitt, M.R., Katz, Y., et al. (2017). A single-cell survey of the small intestinal epithelium. Nature 551, 333–339.

Heijmans, J., van Lidth de Jeude, J.F., Koo, B.K., Rosekrans, S.L., Wielenga, M.C., van de Wetering, M., Ferrante, M., Lee, A.S., Onderwater, J.J., Paton, J.C., et al. (2013). ER stress causes rapid loss of intestinal epithelial stemness through activation of the unfolded protein response. Cell reports 3, 1128–1139.

Hetz, C., and Papa, F.R. (2018). The Unfolded Protein Response and Cell Fate Control. Molecular cell 69, 169–181.

Hollien, J., Lin, J.H., Li, H., Stevens, N., Walter, P., and Weissman, J.S. (2009). Regulated Ire1-dependent decay of messenger RNAs in mammalian cells. The Journal of cell biology 186, 323–331.

Hollien, J., and Weissman, J.S. (2006). Decay of endoplasmic reticulum-localized mRNAs during the unfolded protein response. Science 313, 104–107.

Imagawa, Y., Hosoda, A., Sasaka, S., Tsuru, A., and Kohno, K. (2008). RNase domains determine the functional difference between IRE1alpha and IRE1beta. FEBS letters 582, 656–660.

Iwawaki, T., and Akai, R. (2006). Analysis of the XBP1 splicing mechanism using endoplasmic reticulum stress-indicators. Biochemical and biophysical research communications 350, 709–715.

Iwawaki, T., Hosoda, A., Okuda, T., Kamigori, Y., Nomura-Furuwatari, C., Kimata, Y., Tsuru, A., and Kohno, K. (2001). Translational control by the ER transmembrane kinase/ribonuclease IRE1 under ER stress. Nature cell biology 3, 158–164.

Lee, K., Tirasophon, W., Shen, X., Michalak, M., Prywes, R., Okada, T., Yoshida, H., Mori, K., and Kaufman, R.J. (2002). IRE1-mediated unconventional mRNA splicing and S2P-mediated ATF6 cleavage merge to regulate XBP1 in signaling the unfolded protein response. Genes & development 16, 452–466.

Li, H., Korennykh, A.V., Behrman, S.L., and Walter, P. (2010). Mammalian endoplasmic reticulum stress sensor IRE1 signals by dynamic clustering. Proceedings of the National Academy of Sciences of the United States of America 107, 16113–16118.

Martino, M.B., Jones, L., Brighton, B., Ehre, C., Abdulah, L., Davis, C.W., Ron, D., O’Neal, W.K., and Ribeiro, C.M. (2013). The ER stress transducer IRE1beta is required for airway epithelial mucin production. Mucosal immunology 6, 639–654.

Mi, L.Z., Lu, C., Li, Z., Nishida, N., Walz, T., and Springer, T.A. (2011). Simultaneous visualization of the extracellular and cytoplasmic domains of the epidermal growth factor receptor. Nature structural & molecular biology 18, 984–989.

Miyoshi, H., and Stappenbeck, T.S. (2013). In vitro expansion and genetic modification of gastrointestinal stem cells in spheroid culture. Nature protocols 8, 2471–2482.

Paton, A.W., Beddoe, T., Thorpe, C.M., Whisstock, J.C., Wilce, M.C., Rossjohn, J., Talbot, U.M., and Paton, J.C. (2006). AB5 subtilase cytotoxin inactivates the endoplasmic reticulum chaperone BiP. Nature 443, 548–552.

Paton, A.W., Srimanote, P., Talbot, U.M., Wang, H., and Paton, J.C. (2004). A new family of potent AB(5) cytotoxins produced by Shiga toxigenic Escherichia coli. The Journal of experimental medicine 200, 35–46.

Prischi, F., Nowak, P.R., Carrara, M., and Ali, M.M. (2014). Phosphoregulation of Ire1 RNase splicing activity. Nature communications 5, 3554.

Sepulveda, D., Rojas-Rivera, D., Rodriguez, D.A., Groenendyk, J., Kohler, A., Lebeaupin, C., Ito, S., Urra, H., Carreras-Sureda, A., Hazari, Y., et al. (2018). Interactome Screening Identifies the ER Luminal Chaperone Hsp47 as a Regulator of the Unfolded Protein Response Transducer IRE1alpha. Molecular cell 69, 238-252.e237.

Sundaram, A., Plumb, R., Appathurai, S., and Mariappan, M. (2017). The Sec61 translocon limits IRE1alpha signaling during the unfolded protein response. eLife 6.

Tirasophon, W., Lee, K., Callaghan, B., Welihinda, A., and Kaufman, R.J. (2000). The endoribonuclease activity of mammalian IRE1 autoregulates its mRNA and is required for the unfolded protein response. Genes & development 14, 2725–2736.

Tirasophon, W., Welihinda, A.A., and Kaufman, R.J. (1998). A stress response pathway from the endoplasmic reticulum to the nucleus requires a novel bifunctional protein kinase/endoribonuclease (Ire1p) in mammalian cells. Genes & development 12, 1812–1824.

Tschurtschenthaler, M., Adolph, T.E., Ashcroft, J.W., Niederreiter, L., Bharti, R., Saveljeva, S., Bhattacharyya, J., Flak, M.B., Shih, D.Q., Fuhler, G.M., et al. (2017). Defective ATG16L1-mediated removal of IRE1alpha drives Crohn’s disease-like ileitis. The Journal of experimental medicine 214, 401–422.

Tsuru, A., Fujimoto, N., Takahashi, S., Saito, M., Nakamura, D., Iwano, M., Iwawaki, T., Kadokura, H., Ron, D., and Kohno, K. (2013). Negative feedback by IRE1beta optimizes mucin production in goblet cells. Proceedings of the National Academy of Sciences of the United States of America 110, 2864–2869.

Walter, P., and Ron, D. (2011). The unfolded protein response: from stress pathway to homeostatic regulation. Science 334, 1081–1086.

Wang, X.Z., Harding, H.P., Zhang, Y., Jolicoeur, E.M., Kuroda, M., and Ron, D. (1998). Cloning of mammalian Ire1 reveals diversity in the ER stress responses. The EMBO journal 17, 5708–5717.

Wang, Y., Shen, J., Arenzana, N., Tirasophon, W., Kaufman, R.J., and Prywes, R. (2000). Activation of ATF6 and an ATF6 DNA binding site by the endoplasmic reticulum stress response. The Journal of biological chemistry 275, 27013–27020.

Wielenga, M.C., Colak, S., Heijmans, J., van Lidth de Jeude, J.F., Rodermond, H.M., Paton, J.C., Paton, A.W., Vermeulen, L., Medema, J.P., and van den Brink, G.R. (2015). ER-Stress-Induced Differentiation Sensitizes Colon Cancer Stem Cells to Chemotherapy. Cell reports 13, 490–494.

Wiseman, R.L., Zhang, Y., Lee, K.P., Harding, H.P., Haynes, C.M., Price, J., Sicheri, F., and Ron, D. (2010). Flavonol activation defines an unanticipated ligand-binding site in the kinase-RNase domain of IRE1. Molecular cell 38, 291–304.

Yoshida, H., Matsui, T., Yamamoto, A., Okada, T., and Mori, K. (2001). XBP1 mRNA is induced by ATF6 and spliced by IRE1 in response to ER stress to produce a highly active transcription factor. Cell 107, 881–891.

